# Phase separation of RNA-binding protein promotes polymerase engagement and transcription

**DOI:** 10.1101/2021.03.26.436939

**Authors:** Wen Shao, Xianju Bi, Boyang Gao, Jun Wu, Yixuan Pan, Yafei Yin, Zhimin Liu, Wenhao Zhang, Xu Jiang, Wenlin Ren, Yanhui Xu, Zhongyang Wu, Kaili Wang, Ge Zhan, J. Yuyang Lu, Xue Han, Tong Li, Jianlong Wang, Guohong Li, Haiteng Deng, Bing Li, Xiaohua Shen

## Abstract

An RNA-involved phase-separation model has been proposed for transcription control. Yet, the molecular links that connect RNA binding to the transcription machinery remain missing. Here we find RNA-binding proteins (RBPs) constitute half of the chromatin proteome in embryonic stem cells (ESCs), and some are colocalized with RNA polymerase (Pol) II at promoters and enhancers. Biochemical analyses of representative RBPs—such as PSPC1 and PTBP1—show that the paraspeckle protein PSPC1 not only prevents the RNA-induced premature release of Pol II, and also makes use of RNA as multivalent molecules to promote Pol II engagement and activity, by enhancing the phase separation and subsequent phosphorylation and release of polymerase condensates. In ESCs, auxin-induced acute degradation of PSPC1 leads to genome-wide defects in Pol II phosphorylation and chromatin-binding and nascent transcription. We propose that the synergistic interplay of RBPs and RNA aids in the rate-limiting step of polymerase condensate formation to promote active transcription.

## Main text

Intricate regulation of transcription is central for cell differentiation and function in development^1,2^. Genome-wide studies have revealed the prevalence of pausing of RNA polymerase (Pol) II in promoter-proximal regions of most metazoan genes^3–8^. The activity and release of promoter-paused Pol II into elongation is regulated through the phosphorylation state of an intrinsically disordered C-terminal domain (CTD) of the largest subunit of Pol II^8,9^. Intriguingly, transcription of most active genes occurs in short bursts^10–18^. Imaging-based studies have shown transient residence and clustering in the seconds scale for Pol II that initiates or pauses at the promoter^13,19–26^. It has been estimated that only 1 of 100 Pol II-gene interactions will proceed to productive elongation^13,19^. Dynamic assembly and binding of Pol II during initiation suggests key regulatory events that are necessary to stabilize Pol II binding for transcription elongation.

Transcription is thought to take place at discrete nuclear sites known as transcription ‘factories’, hubs or clusters in the form of phase-separated condensates, which allow efficient compartmentalization and coupling of polymerases engaged at multiple genomic sites^27–35^. Increasing evidence indicates that RNA broadly associates with chromatin and feeds back on transcription and chromatin states^6,36–43^. Recently, it was reported that RNA stimulates transcription factor condensates at low levels but dissolves these condensates at high levels^44,45^. A phase-separation model of RNA-mediated feedback control appears attractive to explain features of transcription processes^45,46^. However, this hypothesis remains inconclusive as the key link that connects RNA to the transcriptional machinery with a characteristic DNA-binding activity is still missing. It is widely believed that eukaryotic transcription is coupled with RNA processing^8,9,47^. RNA-binding proteins (RBPs) constitute a major family of regulators that process and metabolize RNA transcripts from their synthesis to function and to decay^48^. A number of RBPs such as WDR43, DDX21/18/5, SRSF1/2, FUS, hnRNPK/U/L, NCL, and NONO, have been implicated in modulating transcriptional, epigenetic, and signaling responses in various cellular contexts^49–63^. Yet, the direct involvement of RBPs and their interplay with RNA in transcription regulation remain to be proven.

In this study, proteomic profiling reveals abundant and dynamic associations of RBPs with chromatin in ESCs. Surveys of selected RBPs show that they interact with RNA Pol II and preferentially bind regulatory hotspots across the genome, and their knockdown attenuates global transcription. Importantly, through combined *in vitro* biochemical and *in vivo* cellular and systems-level analyses, we delineate the role of PSPC1, a representative RBP, in promoting Pol II engagement and activity during transcription. The synergistic interplay between PSPC1 and RNA in modulating polymerase condensate formation is critically dependent on both the phase-separation and RNA-binding activities of PSPC1, the two biochemical features that are shared by many chromatin-associated RBPs. These results suggest a new angle to reconsider the role of chromatin-bound RNA and its binding proteins in gene regulation beyond the canonical components of the transcription machineries.

### RBPs comprise half of the ESC chromatin proteome

To have a fuller understanding of transcription under the chromatin context, we sought to capture all chromatin-associated proteins, directly or indirectly. We used formaldehyde to crosslink the nuclei isolated from mouse embryonic stem cells (ESCs), precipitated chromatin by ethanol, and then released chromatin proteins by DNase I for mass spectrometry analysis (Extended Data Fig. 1a; Methods). Out of 1,357 chromatin proteins (histones excluded) that were detected to overlap in two biological replicates, 537 proteins are involved in transcription and chromatin-related functions, making up 25% of the protein peptide abundance (Fig. 1a, Extended Data Fig. 1b-c and Supplementary Table 1).

**Fig. 1.**
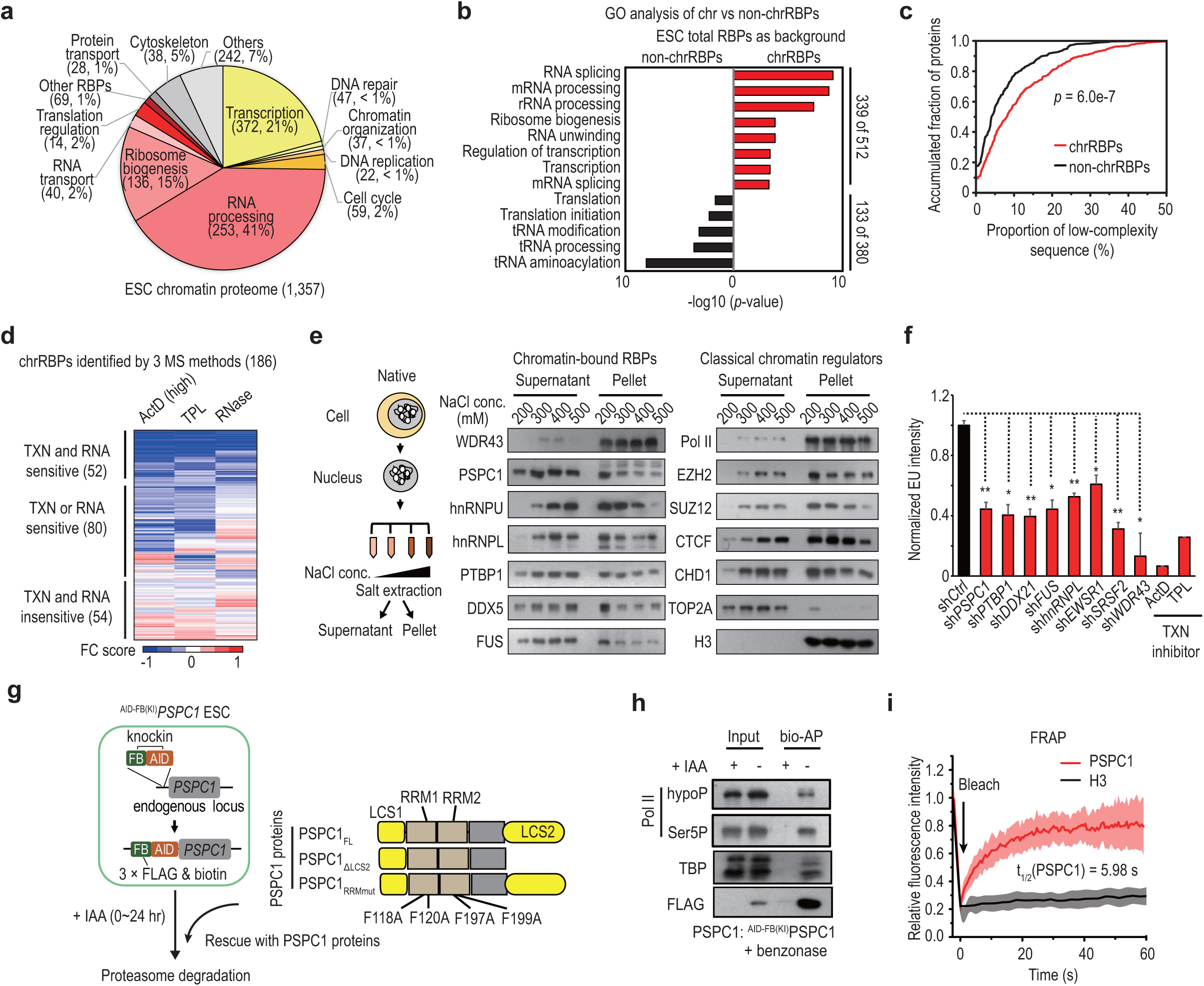
Abundant and dynamic associations of RBPs with chromatin. **a,** Percentages of peptide abundance of chromatin proteins (histones excluded) by intensity-based absolute quantification (iBaq) of mass spec. The protein number and the relative peptide abundance (indicated by iBaq ratio; see Materials and Methods) of functionally associated genes are shown in the brackets. Results are shown as the average value of two independent biological replicates. See also Supplementary Table 1 and Extended Data Fig. 1a-c. **b,** GO analysis of chrRBPs (n = 512) versus non-chrRBPs (n = 380). The total RBPs expressed in ESCs (n = 892) are used as background. The top enriched terms are shown on the y-axis. The x-axis shows enrichment significance by −log10 (*p*-value). Red bars represent terms enriched in chrRBPs; black bars indicate terms enriched in non-chr RBPs. The numbers of RBPs associated with the corresponding terms and the total analyzed RBPs are indicated on the right. **c,** Cumulative distribution curve showing the content of low-complexity sequences in chrRBPs or non-chrRBPs. Other biochemical characterizations (e.g. isoelectric point, intrinsically disordered regions) are shown in Extended Data Fig. 1f. *P*-values, Kolmogorov-Smirnov test. See also Supplementary Table 2. **d,** Heatmap showing the fold change (FC) score of chromatin abundance for the set of 186 chrRBPs identified in all three mass spec (MS) methods. The FC score calculation is described in Materials and Methods. Data are shown as the mean of 4 biological replicates for ActD (high, 1 μg/ml) and RNase, and 3 replicates for TPL. Based on the FC score, chrRBPs are classified into 3 groups with different sensitivities to inhibition of transcription (TXN) and/or RNase treatment (RNA). Representative proteins were listed in Extended Data Fig. 1g. See also Supplementary Table 3. **e,** Protein analysis of native ESC chromatin. Left: pipeline for biochemical extraction by salt in non-crosslinked ESCs; middle and right: western-blot analysis of chrRBPs and classical chromatin regulators. **f,** Quantitative analysis of 5-Ethynyl uridine (EU) incorporation by fluorescence activated cell sorting (FACS) upon depletion of various RBPs in ESCs (see Extended Data Fig. 2a-b). Treatments with transcription inhibitors actinomycin D (ActD) and triptolide (TPL) serve as the positive control. The y-axis shows the average EU intensity normalized to controls cells treated with scramble shRNA (shCtrl). *, *p* < 0.05; **, *p* < 0.01 by two-sided Student’s t-test. **g,** Schematic diagram of the knock-in strategy to construct ^AID-FB(KI)^PSPC1 ESCs and the rescue strategy with wild-type and mutant PSPC1 proteins. **h,** Biotin-mediated affinity purification (bio-AP) of ^AID-FB(KI)^PSPC1 and western-blot analysis. ^AID-FB(KI)^PSPC1 ESCs treated by IAA for 24 hours were used as a negative control. Benzonase was present during cell lysis and bio-AP procedures. **i,** Fluorescence recovery after photobleaching (FRAP) analysis showing the fast recovery of mCherry-PSPC1 puncta in ESCs. See also Extended Data Fig. 2g. GFP tagged-histone H3 was used as a control. The y-axis shows the relative fluorescence intensity normalized to the initial level. Data are shown as mean ± s.d. of 10 biological replicates.

Congruent with previous proteomic analysis^64^, RNA-binding proteins are also significantly enriched (Extended Data Fig. 1d; *p* < 1e-10). By intersecting our chromatin proteome with Tuschl’s RBP repertoire^48^, we defined the overlapping 512 proteins as chromatin-bound RBPs (chrRBPs), which accounts for 62% of the protein abundance on chromatin (Fig. 1a, Extended Data Fig. 1e and Supplementary Table 1). These chrRBPs are enriched in nuclear processes, including RNA processing, splicing, and mRNA transport, in comparison to tRNA and translation-related functions for non-chromatin RBPs (380 proteins) (Fig. 1b, Extended Data Fig. 1e and Supplementary Table 2). Analysis of a published mass spec dataset of proteins pulled down by the CTD of Pol II *in vitro*^65^ revealed that a large proportion (62%, 318) of chrRBPs were detected (>2-fold enrichment) in the CTD interactomes, compared to 32% (123) of non-chromatin RBPs (Supplementary Table 2). In addition, chrRBPs are more positively charged with higher isoelectric points, and intriguingly, exhibit significantly higher contents of low-complexity sequences (LCSs) and intrinsically disordered regions (IDRs) (Fig. 1c, Extended Data Fig. 1f and Supplementary Table 2), which implies a tendency to liquid-liquid phase separation on chromatin^66–70^.

Treatments that inhibit transcription or degrade RNA dramatically attenuated RBP-chromatin associations, but had less effects on transcription factors and epigenetic enzymes (Extended Data Fig. 1a, 1g and 1h, and Supplementary Table 3). Among the 186 chrRBPs that were consistently detected by three quantitative mass spec methods across samples, the majority (71%, 132) exhibited reduced chromatin association in response to at least one treatment (Fig. 1d, Extended Data Fig. 1g, and 1i, and Supplementary Table 3; Methods). Validation of individual proteins showed that 8 tested RBPs fell off the chromatin upon RNA degradation or inhibition of transcription (Extended Data Fig. 1j). PSPC1 and DDX21 appeared to be insensitive to these treatments; however, we could not assume complete RNA degradation by RNases. These results suggest dynamic recruitment of chrRBPs by RNA and/or transcription to chromatin. It also rules out a potential crosslinking artifact. Indeed, we tested 7 chrRBPs under non-crosslinking conditions and found that they all exhibited strong chromatin binding at 200 mM salt in a manner similar to that observed for epigenetic factors (Fig. 1e).

### chrRBPs interact with Pol II and modulate transcription

To explore a potential role of chrRBPs in transcription, we knocked down a number of chrRBPs, including *PSPC1*, *PTBP1*, *DDX21, FUS*, *HNRNPL*, and *EWSR1*. Their depletion caused global reduction of nascent transcripts that were pulse-labeled by 5-ethynyl uridine (EU) (Fig. 1f and Extended Data Fig. 2a-b). To test their interactions with Pol II, we performed co-immunoprecipitation (co-IP) in native ESC lysates treated with benzonase which degrades RNA/DNA. Pol II in various phosphorylation states captured all 8 chrRBPs tested, including PSPC1, PTBP1, DDX21, FUS, and HNRNPL, with different specificity (Extended Data Fig. 2c and 2d). We reported previously that the paraspeckle protein PSPC1 regulates the expression of retroviral ERVL and ERVL-associated genes by promoting TET2-chromatin occupancy in ESCs^58^. Because of its strong binding to chromatin, we then chose PSPC1 as a representative RBP for in-depth characterization.

To efficiently capture endogenous PSPC1 and manipulate its protein levels in a cell, we constructed homozygous knock-in ESCs that carry an in-frame insertion of FLAG and biotin tags fused with an auxin-inducible degron (AID) epitope inserted into the 5’ end of the endogenous *PSPC1* alleles (referred to as ^AID-FB(KI)^PSPC1; Fig. 1g). With this cellular platform, we could simultaneously tag and degrade the endogenous PSPC1 protein. Congruent with the front results, endogenously tagged ^AID-FB(KI)^PSPC1 captured initiating and paused Pol II, represented by hypo-phosphorylated (hypoP) and phosphorylated at serine 5 of the CTD (Ser5P), respectively (Fig. 1h). PSPC1 co-IP also captured TATA-box binding protein (TBP), the first protein that binds to DNA to initiate assemblage of the preinitiation complex (PIC) and Pol II^2,71^. Immunofluorescence analysis showed that PSPC1 exhibited punctate signals that partially overlapped with Pol II and TBP puncta (Extended Data Fig. 2e and 2f). Particularly, PSPC1 nuclear foci exhibited a fast fluorescence recovery (∼5.98 seconds) compared to histone H3 (∼100 seconds) after photobleaching (Fig. 1i and Extended Data Fig. 2g), and were dissolved by inhibition of weak hydrophobic interactions by 1,6-hexanediol (Extended Data Fig. 2h), suggesting the properties of liquid-like condensates.

### PSPC1 promotes CTD incorporation, phosphorylation, and release

PSPC1 contains two LCSs with a 66% IDR content. The LCS2 at the carboxyl terminus of PSPC1 is relatively large with ∼200 residues in length and enriched in hydrophobic glycine (G) and proline (P) residues (Fig. 1g and Extended Data Fig. 3a). TBP comprises an IDR enriched in glutamine (Q) at its amino terminus (51% IDR content; Extended Data Fig. 3a). Indeed, recombinant full-length PSPC1_FL_ protein was able to form spherical liquid-like droplets at around its estimated nuclear concentrations (5 μM) (Extended Data Fig. 3b-d and Supplementary Table 1; Methods). In comparison, recombinant TBP protein formed fiber-like irregular aggregates in the absence of dextran, but was able to form liquid-like droplets in the presence of dextran, at a concentration of 5 μM, which is well above its estimated nuclear concentration of 0.06∼0.3 μM (Extended Data Fig. 3a-c, 3e, and Supplementary Table 1; Methods). Given the well-recognized role of TBP in transcription initiation, we regarded TBP droplets as a surrogate for the more complex *in vivo* initiation condensates.

Both PSPC1 and TBP droplets incorporated recombinant CTD (with 20 heptad repeats) inside at a CTD concentration of 0.6 μM which is around the estimated nuclear concentration of Pol II, while CTD failed to phase separate on its own (Extended Data Fig. 3c-e). Addition of PSPC1_FL_ to the TBP and CTD mix produced bigger and brighter CTD droplets, which exhibited liquid-like fusion behaviors and were quickly dissolved by 1,6-hexanediol (Fig. 2a-b and Extended Data Fig. 3f-h). Droplet sedimentation analysis also confirmed that ∼3-fold more CTD protein was trapped inside PSPC1_FL_-TBP-CTD droplets compared with TBP-CTDdroplets (Fig. 2c and Extended Data Fig. 3i). By contrast, the PSPC1_ΔLCS2_ mutant that lacks the LCS2 poorly phase-separated on its own, and failed to affect formation of TBP-CTD droplets (Fig. 1g, 2a-c, and Extended Data Fig. 3b, 3f and 3i). In comparison, an RNA-binding mutant of PSPC1, designated as PSPC1_RRMmut_ that carries four point mutations (F118A, F120A, K197A and F199A) in the RRMs^58^, was able to form liquid droplets and promoted formation of TBP-CTD droplets in a degree weaker than PSPC1_FL_ (Fig. 2a-c and Extended Data Fig. 3f and 3i). These results indicate that the LCS-mediated phase-separation activity of PSPC1 promotes the CTD incorporation.

**Fig. 2.**
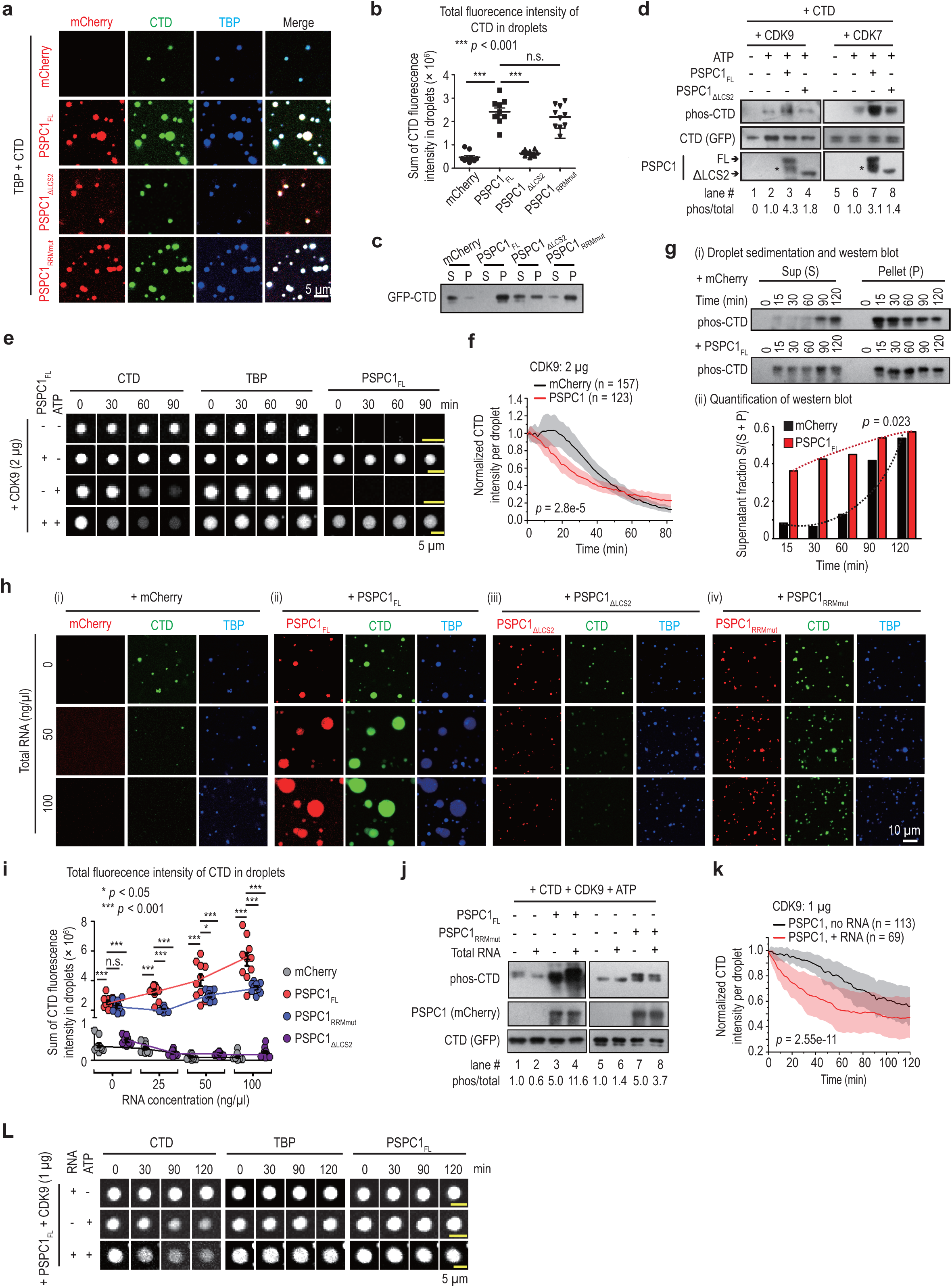
RNA synergizes the effects of PSPC1 in promoting the incorporation, phosphorylation, and release of the CTD via phase separation. **a-c,** Droplet formation assay of TBP (5 μM) and CTD (0.6 μM) with full-length (FL) or mutant PSPC1 proteins (5 μM) or mCherry (5 μM). The calculation of protein’s physiological concentration is described in Materials and Methods. The same protein concentrations and experimental conditions were used in all droplet assays of this paper unless otherwise indicated. All the samples were acquired at the same time. Representative pictures are shown in panel **a**. Panel **b,** shows the quantification of total fluorescence intensity of CTD in droplets shown in panel **a**,. The y-axis shows the sum of fluorescence intensity of CTD in droplets in each field of view. N = 10 fields for each condition. *P*-values, two-sided Student’s t-test. n.s., not significant. The quantification of droplet sizes is shown in Extended Data Fig. 3f. **c,** Droplet sedimentation and western-blot analysis. Supernatant (S) represents the free proteins. Pellet (P) represents proteins inside droplets. The schematic diagram and quantification of western-blot are shown in Extended Data Fig. 3i. **d,** Kinase assay showing that PSPC1_FL_ but not PSPC1_ΔLCS2_ promotes CTD phosphorylation. About 5 μg of mCherry (lane # 1, 2, 5 and 6) or PSPC1_FL_ (lane # 3 and 7) or PSPC1_ΔLCS2_ (lane # 4 and 8) were incubated with recombinant CTD protein (0.2 μg) and CDK9/CDK7 (0.2 μg) in the presence or absence of ATP (0.1 mM) as indicated. The quantified ratio of phosphorylated (phos) versus total level of the CTD by western blots (middle) are shown at the bottom. We note that recombinant PSPC1_FL_ protein is prone to fragmentation during protein purification. The asterisk indicates the fragmented protein. See also Extended Data Fig. 4a. **e-g,** Time-lapse analysis of CTD release by imaging (**e** and **f**) and by sedimentation assays **(g)**. To initiate the release reaction, CDK9 (2 μg) and ATP (final 0.1 mM) were added to pre-assembled droplets with indicated components. Panel **e,** shows images taken from representative droplets under each condition. The CTD channel (left), TBP channel (middle) and PSPC1_FL_ channel (right) of individual droplets were recorded simultaneously. See also Extended Data Fig. 4c and Supplementary Video 1 and 3. Quantification of CTD intensity **(f)**, was based on images shown in panel **e** (see also Extended Data Fig. 4b and Supplementary Video 1). The y-axis shows the relative CTD intensity of individual droplet normalized to its TBP intensity (see Methods). Data are shown as mean ± s.d. of 157 droplets for the mCherry group and 123 droplets for the PSPC1 group. *P*-value, Kolmogorov-Smirnov test. The time-course quantification of relative CTD release rate is shown in Extended Data Fig. 4b. Consistent with previous reports^77^, addition of CDK9 in the presence of ATP led to a gradual loss of CTD fluorescence signals with time from TBP condensates, indicating of a phosphorylation-dependent release. By comparison, phosphorylation did not affect the phase separation of TBP and PSPC1_FL_. Panel **g**, shows time-lapse sedimentation analysis of CTD release in another independent experiment. At each indicated time point, droplets and free proteins were collected by sedimentation for western-blot analysis (i),. The supernatant fraction S/(S + P) was calculated from quantification of the western blots (ii). The comparison of the absolute level of phos-CTD in the supernatant was shown in Extended Data Fig. 4D. *P*-value, two-tailed Student’s paired t-test for the comparison of supernatant fraction between two groups at each time point. **h-i,** Effects of RNA and PSPC1 on CTD incorporation into TBP droplets. Panel **h,** shows representative pictures of phase-separated TBP-CTD droplets with the addition of mCherry (i), PSPC1_FL_ (ii), PSPC1 _Δ LCD2_ (iii), PSPC1_RRMmut_ (iv) in the presence of 0-100 ng/μl of total RNA from ESCs. Panel **i** shows quantifications of total fluorescence intensity of CTD in droplets. N = 10 fields for each condition. *P*-values, two-sided Student’s t-test. n.s., not significant. Quantification of droplet size and total fluorescence intensity of TBP in droplets are shown in Extended Data Fig. 4e-f. **j,** Effects of RNA and PSPC1 on CTD phosphorylation. The kinase assay was performed under the same condition as in panel **d**. **k-l,** Time-lapse imaging analysis of CTD release with or without RNA (50 ng/μl). Addition of RNA led to rapid decreases of CTD signals in the PSPC1-TBP droplets upon the onset of imaging inquiry. In order to dissect the effect of RNA, we slowed down the time course of CTD release by adding half amount of CDK9 enzyme (1 μg) compared to that used (2 μg) in panels **e-f** to slow down the kinase reaction. Panel **k** shows the time-course quantification of relative CTD intensity of individual droplet shown in panel **l**. The y-axis is the normalized CTD intensity (see Methods). Data are shown as mean ± s.d. of 113 droplets for the ‘PSPC1, no RNA group’ and 69 droplets for the ‘PSPC1, + RNA’ group. *P*-value, Kolmogorov-Smirnov test. The time-course quantification of relative CTD release rate is shown in Extended Data Fig. 4h. Panel (**l**) shows images taken from representative droplets under each condition. The CTD channel (left), TBP channel (middle) and PSPC1_FL_ channel (right) were recorded simultaneously for individual droplets. See also Extended Data Fig. 4i and Supplementary Video 2 and 3. In panels **e, g** and **j**, we used anti-Pol II Ser5P antibody to detect phos-CTD. In panels **h-l**, total RNA from ESCs was used.

Hyper-phosphorylation of the Pol II CTD is required for its activity and release in cells^8,72^. Next, we tested the effects of PSPC1 on the phosphorylation and release of CTD in the presence of recombinant CTD kinases CDK7 or CDK9 *in vitro*. PSPC1_FL_ protein markedly enhanced CTD phosphorylation in a PSPC1_FL_ dose-dependent manner, whereas PSPC1_ΔLCS2_ and the control bovine serum albumin (BSA) had no effect (Fig. 2d and Extended Data Fig. 4a). In accordance with increased CTD phosphorylation, PSPC1_FL_ led to a more rapid release of CTD from TBP-PSPC1_FL_ droplets compared to droplets containing TBP alone (Fig. 2e, rows 3 and 4; Fig. 2f and Extended Data Fig. 4b-c; Supplementary Video 1). PSPC1_FL_ skewed the release rate curve to early time points from a peak time at ∼37 minutes to ∼10 minutes following addition of ATP (Extended Data Fig. 4b; Methods). Droplet sedimentation analysis also confirmed an accelerated release of phosphorylated CTD to 15 minutes—the earliest time point analyzed (Fig. 2g and Extended Data Fig. 4d). We noted that CDK9-mediated phosphorylation did not affect the phase separation of TBP and/or PSPC1 (Fig. 2e and Extended Data Fig. 4c; Supplementary Video 1). Taken together, PSPC1 not only promotes the incorporation of unphosphorylated CTD into TBP initiation droplets, but also accelerates CDKs-mediated phosphorylation and release of CTD through phase separation.

### PSPC1 prevents RNA-induced CTD eviction and synergizes with RNA to promote CTD incorporation, phosphorylation, and release

Phase diagram and droplet sedimentation showed that addition of total RNA from ESCs gradually promoted the formation of PSPC1 droplets in a PSPC1 and RNA concentration-dependent manner (Extended Data Fig. 3d and 3j). High RNA levels (up to 100 ng/μl tested) led to smaller droplets and the appearance of irregular fiber-like aggregates (Extended Data Fig. 3d). Compared to TBP alone, TBP-PSPC1 droplets were also dramatically increased by RNA (up to 100 ng/μl RNA) (Fig. 2h, panel ii; Extended Data Fig. 4e-f, red color). These enhancement effects of RNA were impaired when substituted with PSPC1_ΔLCS2_ and PSPC1_RRMmut_ mutants (Fig. 2h, panel iii and iv; Extended Data Fig. 4e-f). Note that PSPC1_RRMmut_ and TBP with minimal or no RNA-binding activity appeared to be less sensitive to RNA (Extended Data Fig. 3e; Extended Data Fig. 4f, gray color; Extended Data Fig. 4g). Such minimal changes may result from nonspecific electrostatic effect between RNA and protein. Thus, RNA acts as a multivalent ligand to promote PSPC1 phase behavior within the range of balanced RNA:protein interactions, while high RNA levels that disrupt this balance may suppress liquid-liquid phase separation via gelation or dissolve the phase via repulsive-like charge interactions.

Next, we tested the effects of RNA on the condensate-interacting behaviors of the CTD. Interestingly, in the absence of PSPC1, RNA led to a gradual loss of CTD fluorescence from TBP droplets in an RNA dosage-dependent manner (Fig. 2h, panel i; Fig. 2i, grey color), while TBP droplets were yet to be formed (Extended Data Fig. 4f, grey). This effect was independent of RNA sequences tested (data not shown), suggesting that imbalanced negative charge interactions evict CTD, mimicking phosphorylation’s effect on CTD. This RNA-induced CTD eviction was completely blocked by the addition of PSPC1_FL_, which further increased the CTD incorporation into TBP droplets (Fig. 2h, panel ii; Fig. 2i and Extended Data Fig. 4e, red color). In contrast, PSPC1_ΔLCS2_ failed to block CTD eviction by RNA (Fig. 2h, panel iii; Fig. 2i and Extended Data Fig. 4e, purple color). Although PSPC1_RRMmut_ prevented RNA-induced eviction of CTD, TBP-PSPC1_RRMmut_ droplets remained small in size and the levels of incorporated CTD did not scale with RNA concentrations (Fig. 2h, panel iv; Fig. 2i and Extended Data Fig. 4e, blue color).

Simultaneous addition of PSPC1_FL_ and RNA dramatically enhanced CDK9-mediated phosphorylation of the CTD by 12∼19-fold, compared to a 5-8-fold increase by PSPC1_FL_ alone, whereas RNA alone showed no obvious effect (Fig. 2j, lanes 1-4). By comparison, PSPC1_RRMmut_ led to a moderate increase (2.6∼5.0-fold) regardless of addition of RNA (Fig. 2j, lanes 5-8). Moreover, RNA synergized with PSPC1_FL_ in enhancing the release of phosphorylated CTD only when ATP was present (Fig. 2k-l and Extended Data Fig. 4h-i; Supplementary Video 2 and 3). Taken these results together, PSPC1 not only neutralizes the effect of RNA to expel CTD, and also makes use of RNA to efficiently compartmentalize CTD for enhanced phosphorylation and release in the presence of CDKs. The synergistic interplay between PSPC1 and RNA is critically dependent on the phase-separation and RNA-binding activities of PSPC1 (Extended Data Fig. 4j).

### PSPC1 stabilizes Pol II engagement during *in vitro* transcription

Next, we examined the effect of PSPC1 on the Pol II enzyme in a fully defined *in vitro* transcription system. We utilized a DNA template (Extended Data Fig. 5a) containing a heteroduplex bubble which has been widely used as a nucleic acid scaffold in Pol II structural studies^73^. Pol II can bind to the single-stranded DNA within the bubble without the help of general transcription factors (Fig. 3a and Extended Data Fig. 5a-c). Addition of di-nucleotide UpG guide RNA and nucleoside triphosphates (NTPs) facilitated Pol II elongation to generate a 278-nt full-length run-off transcript (Fig. 3b, lane 5). When GTP was omitted (NTPs-GTP), Pol II produced a 33-nt short G-less transcript then stalled at the triple-C template site (Fig. 3b, lanes 1 and 3).

**Fig. 3.**
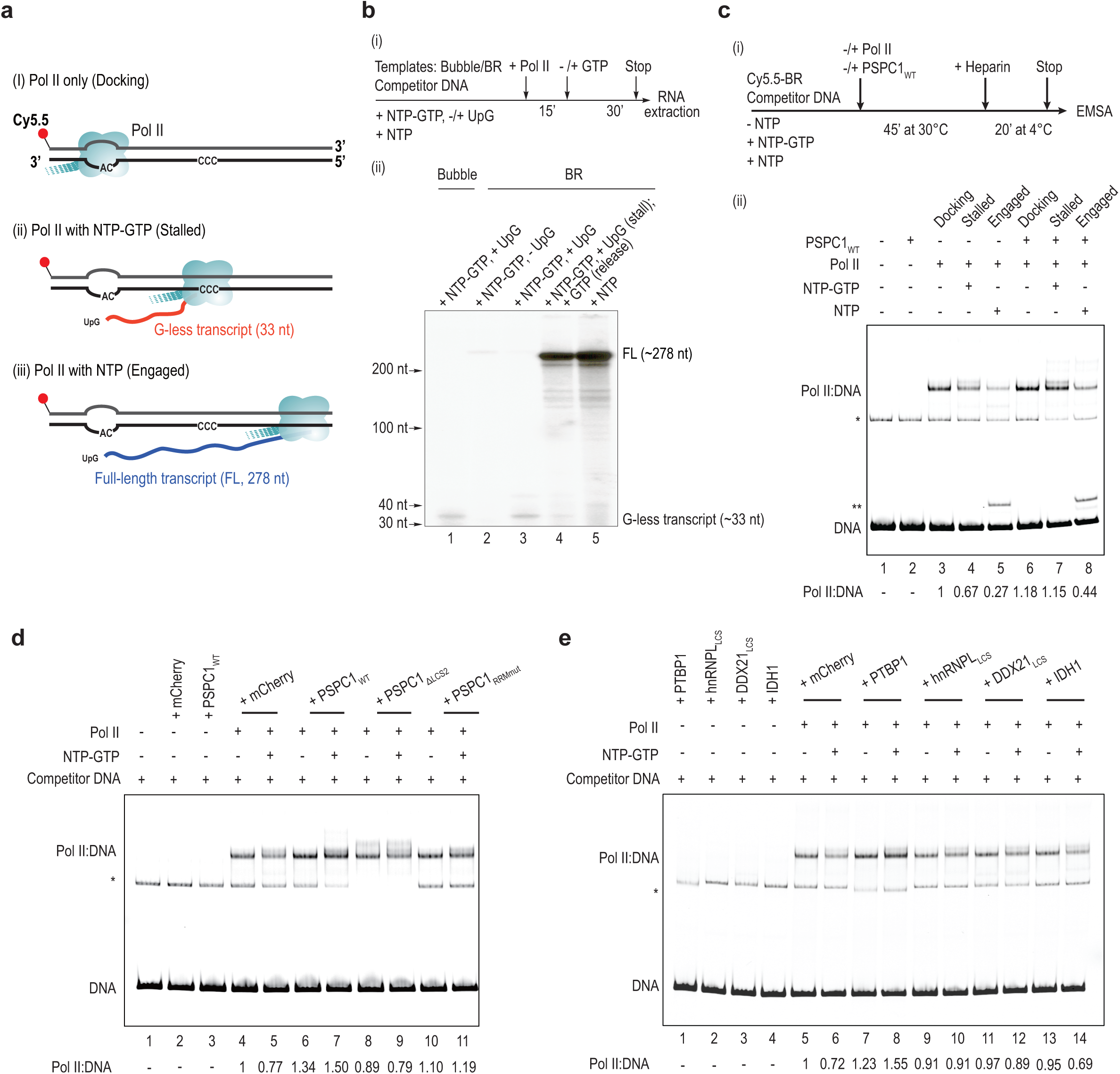
PSPC1 stabilizes Pol II engagement during transcription *in vitro*. **a,** Schematic diagram of the *in vitro* transcription system. In the absence of NTPs, Pol II binds to the bubble structure on the template without transcription (i). With the addition of NTPs-GTP and UpG, Pol II initiates transcription at the AC site and pauses at the CCC site on the template, producing a short 33-nt transcript (ii). In the presence of NTPs, Pol II runs off the template and produces a ∼278-nt transcript. See also Extended Data Fig. 5a. **b,** RNA detection of *in vitro* transcription. Bubble only and bubble coupled with the right 601 sequence (BR) were used as the templates as indicated (see Extended Data Fig. 5a). Pol II and NTPs were added to initiate transcription (i), 32P-labeled CTP was used for detection (ii). G-less transcripts (∼33 nt) and full-length transcripts (∼278 nt) were shown as indicated. **c-e,** EMSA of Pol II and BR template during *in vitro* transcription. The BR template (Cy5.5-labeled), Pol II, and PSPC1 or mCherry control proteins in the absence or presence of NTPs as indicated were incubated at 30 °C for 45 minutes. Heparin or mock was added and incubated at 4 °C for additional 20 minutes to remove unengaged Pol II (see also Extended Data Fig. 5c). The free template (DNA) and the supershifted Pol II:DNA bands are indicated. The bands marked by single asterisk are likely to be a nonspecific byproduct from gel purification during the bubble-template assembly. The bands marked by double asterisks is likely to be a DNA:RNA hybrid (R-loop), given its sensitivity to RNase H (data not shown). The relative Pol II:DNA binding intensity was quantified and indicated at the bottom of each data figure. See also Extended Data Fig. 5f and 5i.

We then performed an electrophoretic mobility shift assay (EMSA) to measure Pol II binding to the template by quantifying the supershifted Pol II:DNA signal. To minimize loosely docked Pol II, we added heparin, which competes with the template to occupy the DNA-binding channel of Pol II (Extended Data Fig. 5c). Heparin reduced Pol II binding to the template in the absence of NTPs, but had negligible effects on the stalled or elongating Pol II (Extended Data Fig. 5c; Extended Data Fig. 5d, lane 5 vs lane 11). Transcription led to a gradual decrease of supershifted Pol II:DNA signals, for both the stalled Pol II (+ NTPs-GFP) and the engaged Pol II (+ NTPs) (Fig. 3c). This observation is consistent with the nuclear transcription whereby Pol II frequently falls off the chromatin template during initiation and promoter pausing^13,19^.

To test the effect of PSPC1 on Pol II engagement in this *in vitro* transcription system, we first titrated a double-stranded DNA competitor to prevent nonspecific binding of PSPC1 to the bubble DNA template (Extended Data Fig. 5e). PSPC1 did not form a stable complex with Pol II, suggesting weak interactions. Interestingly, addition of PSPC1_FL_ consistently enhanced the Pol II:DNA signals in the absence or presence of NTPs or NTPs-GTP (Fig. 3c, lanes 3-5 vs 6-8; Extended Data Fig. 5d and 5f), whereas PSPC1_RRMmut_ and PSPC1_ΔLCS2_ had negligible effects (Fig. 3d, lanes 8-11; Extended Data Fig. 5f-g). These results indicate that PSPC1 directly promotes the Pol II-DNA engagement during the initial loading and subsequent pausing and elongation stages.

To test whether this enhancement effect is specific to PSPC1, we tested several recombinant proteins, including Polypyrimidine Tract Binding Protein 1 (PTBP1), the truncated LCS domains of hnRNPL (hnRNPL_LCS_) and DDX21 (DDX21_LCS_), and isocitrate dehydrogenase IDH1. It is known that PTBP1 binds to polypyrimidine tract of pre-mRNA introns^74^, and IDH1 binds directly to GA- or AU-rich RNA^42^. PTBP1 (41% IDR), hnRNPL_LCS_, and DDX21_LCS_, but not IDH1(14% IDR), were able to phase separate and incorporate CTD inside their droplets in the presence of dextran (Extended Data Fig. 5h and data not shown). Only the addition of PTBP1, but not the other recombinant proteins tested, increased Pol II:DNA supershifted signals (Fig. 3e and Extended Data Fig. 5i). The stimulatory effect of PSPC1 and PTBP1 implies that many chrRBPs could act similarly to promote Pol II engagement during transcription. As for PSPC1 mutants, IDH1, hnRNPL_LCS_, and DDX21_LCS_ proteins which do not have the capability to bind RNA and phase separate at the same time, they all failed to show an obvious effect. These results demonstrate that both the RNA-binding and phase-separation activities are necessary for an RBP to promote Pol II engagement during *in vitro* transcription.

### PSPC1 co-localizes with Pol II and its acute degradation impairs transcription

We then sought to explore the *in vivo* function of PSPC1 in regulating Pol II transcription. We first mapped its chromatin-binding sites by chromatin immunoprecipitation following by sequencing (ChIP-seq). The overall targets of endogenously and ectopically tagged PSPC1 are highly similar (*p* < 2.2e-16 by Fisher’s exact test) and overlap extensively with those of initiating (hypoP) and paused (Ser5P) Pol II (Extended Data Fig. 6a-d and Supplementary Table 4; Methods). Among a total of 11,589 overlapping PSPC1 peaks, 53% are localized in the promoters of 5,262 genes, and 6.1% are in enhancers (Fig. 4a, Extended Data Fig. 6e and Supplementary Table 4). PSPC1 binding is also enriched at the TSS mimicking that of hypoP Pol II, and is positively correlated with active histone marks and gene expression (Fig. 4b and Extended Data Fig. 6f-g). In addition, treatments of ESCs with 1,6-hexanediol, which perturbs weak hydrophobic interactions, abolished PSPC1 binding to its target genes (Extended Data Fig. 6h). Moreover, PSPC1_ΔLCS2_ and PSPC1_RRMmut_ mutants exhibited significantly reduced binding at the genome-wide level and in individual genes, and showed diffused nuclear distributions in contrast to punctate staining of PSPC1_FL_ (Fig. 4c-d and Extended Data Fig. 6i-k). These results indicate that both phase-separation and RNA-binding activities of PSPC1 are required for its efficient targeting to chromatin.

**Fig. 4.**
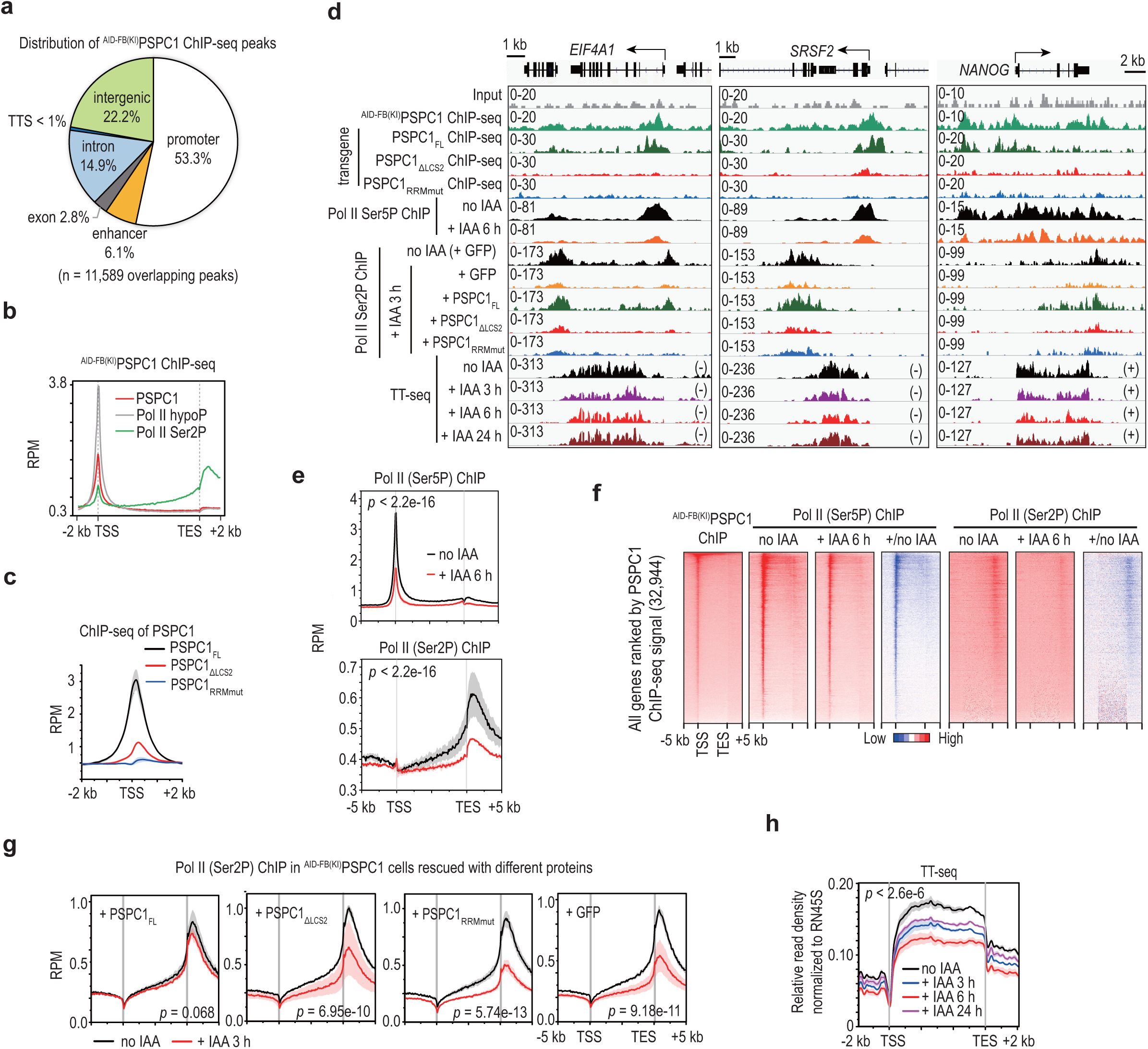
PSPC1 promotes Pol II binding and nascent transcription in ESCs. **a-b,** ChIP-seq analysis of ^AID-FB(KI)^PSPC1. Panel **a,** shows the distribution of ^AID-FB(KI)^PSPC1 ChIP-seq peaks. Among a total of 11,589 overlapping peaks in two biological replicates, 53% are localized in the promoters of 5,262 genes, and 6.1% are in enhancers and super-enhancers (see also Extended Data Fig. 6e and Supplementary Table 5). Panel **b,** shows metagene analysis of PSPC1 and Pol II ChIP-seq signals across all mouse genes (n = 32,944). The y-axis shows reads per million reads (RPM). **c,** ChIP-seq analysis of PSPC1 transiently expressed in ESCs. The wild-type and mutant proteins of PSPC1 are shown in Fig. 1g. **d,** UCSC genome browser view of ChIP-seq and TT-seq at representative loci. This data figure includes ChIP-seq tracks for ^AID-FB(KI)^PSPC1 (Panel **a**), Pol II (Panels **b** and **e-g**), and transiently expressed PSPC1 proteins (Panel **c**). **e-f,** ChIP-seq analysis of Pol II Ser5P and Ser2P upon PSPC1 degradation induced by IAA for 6 hours. Panel **e,** shows metagene analysis across all mouse genes (n = 32,944). Panel **f,** shows heatmaps of PSPC1 and Pol II ChIP-seq signals, and the ratio of changes of Pol II ChIP-seq signals before and after adding IAA (6 hours). The heatmaps are sorted by PSPC1 ChIP-seq signal on the left. See also Extended Data Fig. 6m-n. **g,** Metagene analysis of Pol II Ser2P ChIP-seq in ^AID-FB(KI)^PSPC1 ESCs that were transiently transfected with the wild-type and mutant PSPC1 proteins in the presence or absence of IAA (3 h). Transfection with the *GFP* plasmid serves as the negative control. Also see Pol II Ser2P tracks in the lower middle of panel **d**. **h,** Metagene analysis of transient transcriptome sequencing of nascent transcripts (TT-seq) during the time course of PSPC1 degradation induced by IAA. The y-axis shows the relative read density of nascent transcripts across all protein-coding genes (n = 20,516) normalized to the *RN45S* rRNA. Also see TT-seq tracks in the panel **d**. In panels **c**, **e**, **g** and **h**, data are shown as mean ± s.d. of 2 biological replicates. *P*-values, Kolmogorov-Smirnov test. In panels **b**, **c**, **e** and **g**, the y-axis shows reads per million reads. (RPM). In panels **b, c, e, f** and **g**, metagene and heatmap analyses were plotted around the gene body or TSS of all mouse genes (n = 32,944).

We then examined the primary effects of PSPC1 degradation at a time scale that preclude indirect consequences using ^AID-FB(KI)^PSPC1 ESCs. Addition of the auxin analog indole-3-acetic acid (IAA) induced rapid degradation of PSPC1 protein, which was reduced to <40% at 1 hour and became barely detectable at 4 hours (Extended Data Fig. 6l). The protein levels of phosphorylated Pol II, but not total Pol II, were dramatically decreased to 20-30% at 4 hours (Extended Data Fig. 6l). Levels of Pol II phosphorylation recovered after initial decreases during prolonged treatment of IAA, which implies compensatory mechanism(s) that safeguard the steady-state Pol II activity. Consistently, ChIP-seq showed reduced binding of Ser5P Pol II at the TSS and elongating Pol II (phosphorylation at serine 2 of the CTD, Ser2P) across gene-bodies and downstream regions at 3 and 6 hours of IAA treatment (Fig. 4e and Extended Data Fig. 6m). The degree of downregulation in Pol II ChIP signals was positively correlated with the PSPC1 ChIP signal (Fig. 4f and Extended Data Fig. 6n). Importantly, transient expression of the full-length PSPC1_FL_, but not PSPC1_ΔLCS2_ or PSPC1_RRMmut_ mutant, rescued the genome-wide reduction in Ser2P Pol II binding to chromatin (Fig. 4g). This indicates that PSPC1 utilizes its phase separation and RNA-binding activity to stabilize Pol II binding *in vivo*.

Transient transcriptome sequencing (TT-seq) of nascent transcripts further revealed downregulated transcription that occurred at an early time point of 3 hours after adding IAA. TT-seq signals became the lowest at 6 hours and remained lower than the level prior to PSPC1 degradation despite a slight recovery at 24 hours (Fig. 4h and Extended Data Fig. 6o). Thus, global transcriptional reduction corresponds with the early defects in phosphorylation and chromatin binding of Pol II upon degradation of PSPC1, demonstrating a direct role for PSPC1 in regulating Pol II transcription dynamics *in vivo*. Of note, paraspeckles are absent in ESCs, as they lack expression of the long isoform of the structural noncoding RNA *Neat1*^75^. Therefore, the observed functions of PSPC1 in transcription are independent of its previously known role as paraspeckle components.

### Genome-wide colocalization of chrRBPs and Pol II correlates with active transcription

To explore a general role for chrRBPs in transcription, we had a glance at where they bind in the genome. We performed ChIP-seq in ESCs for RNA chaperone hnRNPU, the nuclear matrix proteins SAFB1 and SAFB2, and the proteins UTP3, UTP6, and CIRH1A that are known as components of the small-subunit processome (SSUP). We also re-analyzed 7 published ChIP-seq datasets (WDR43, hnRNPK, SRSF2, NONO, DDX21, LIN28A, and METTL3; Supplementary Table 4). Similar to what we have observed for PSPC1, all analyzed chrRBPs bind strongly to regulatory DNA elements, including TSSs, enhancers, and super-enhancers (Fig. 5a-b and Extended Data Fig. 7a), in line with the previous observation in human HepG2 and K562 cells^60^.

**Fig. 5.**
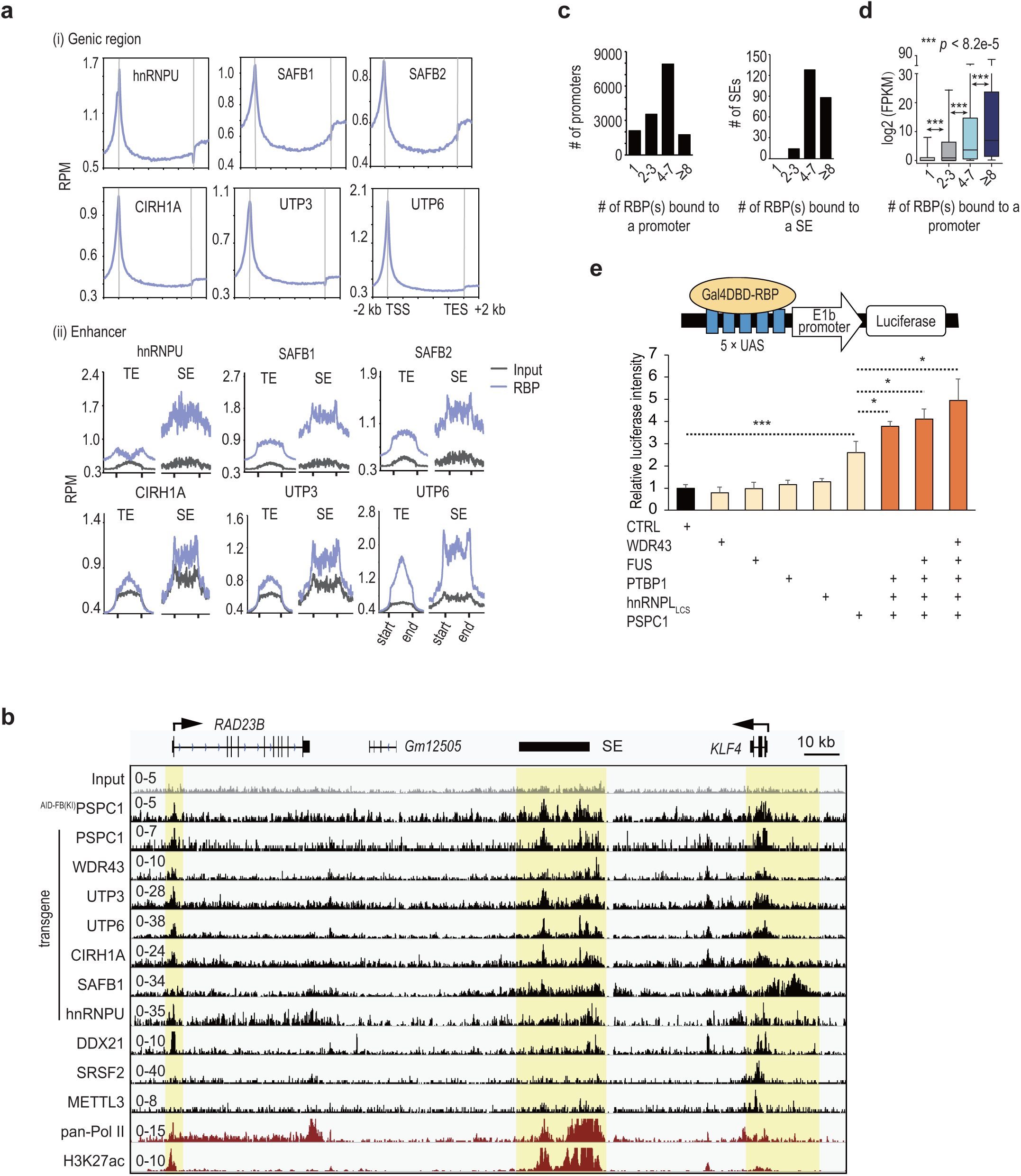
Co-localization of RBPs at promoters and enhancers modulates transcription. **a,** Metagene analysis of ChIP-seq signals of various RBPs across all mouse genes (n = 32,944) (i), and enhancers (ii). ChIP-seq was performed in ESCs. The y-axis is reads per million reads (RPM). TE, typical enhancers (n = 8,704). SE, super enhancers (n = 231). Results of other RBPs analyzed are shown in Extended Data Fig. 7a. Similar to the reported role of another SSUP component WDR43^61^, UTP3, UTP6 and CIRH1A also bind prevalently to active gene promoters, which suggests moonlight functions of SSUP in coordinating Pol I and Pol II transcription. **b,** UCSC genome browser view of ChIP-seq tracks for selected RBPs, pan-Pol II and H3K27ac at representative loci. **c,** Extensive co-localization of RBPs at enhancers and promoters. Gene promoters or super enhancers (SEs) were divided into 4 groups based on the numbers of bound RBPs. The y-axis shows the number of promoters (left) and super enhancers (right) bound by different numbers of RBPs. See also Supplementary Table 4. **d,** Boxplot showing a positive correlation between the number of co-bound RBPs (x-axis) and gene expression (y-axis) *P*-values, two-sided Student’s t-test. **e,** Promoter tethering assay in 293T cells. Tethering of PSPC1 to the reporter gene promoter enhanced luciferase expression by ∼2.5-fold. Despite negligible effects shown by hnRNPL_LCS_, PTBP1, FUS, and WDR43 individually, simultaneous tethering of PSPC1 with these RBPs led to an incremental increase of luciferase activity as a function of the number of RBPs co-tethered. A maximum of ∼5-fold enhancement was observed when all five RBPs were co-expressed. For each assay, the same amount of RBP fused with Gal4 DNA-binding domain (DBD) was co-transfected. The y-axis shows the relative luciferase intensity normalized to the Gal4DBD control. Data are shown as mean ± s.d. of ≥ 3 biological replicates. *, *p* < 0.05; ***, *p* < 0.001 by two-sided Student’s t-test.

This set of 14 chrRBPs intensively co-occupy a total of 15,317 promoters and 231 super-enhancers, of which 77% (11,730) and 92% (212), respectively, are also targeted by RNA Pol II (Fig. 5c and Supplementary Table 5). Remarkably, ∼1,376 promoters are co-bound by ≥8 chrRBPs, and ∼8,234 by 4-7 chrRBPs, and ∼13,191 (86%) are bound by ≥2 chrRBPs (Fig. 5c). Over 98% of super-enhancers (∼226) are co-bound by ≥3 chrRBPs. The degree of co-binding positively correlates with the level of mRNA expression (Fig. 5d). Unsupervised clustering also revealed a strong positive correlation with Pol II and active histone marks and transcription regulators such as MED1, OCT4, and NANOG, but relatively poor-correlation with repressive marks (Extended Data Fig. 7b).

Consistent with the genome-wide colocalization of multiple RBPs, simultaneous addition of PSPC1, PTBP1, and hnRNPL_LCS_ produced larger droplets and incorporated more CTD than single RBPs (Extended Data Fig. 7c-d). In addition, tethering of PSPC1 alone or together with PTBP1, hnRNPL_LCS_, FUS, and WDR43 to a synthetic promoter led to 2.5-5-fold incremental increases of luciferase activity, a functional correspondence to the number of proteins co-tethered (Fig. 5e). These results imply that diverse RBPs might act collaboratively to form transcription condensates to enhance polymerase incorporation and activity. This notion suggests a functional explanation for the prevalent co-binding of RBPs at the regulatory hotspots of the genome.

## DISCUSSION

Here we reveal that hundreds of RBPs are dynamically present on chromatin with their numbers and abundance surpassing even those of classic epigenetic and transcription factors. Surveys of selected RBPs show that they tend to interact and colocalize with Pol II at the genome-wide level, and their knockdown attenuates and co-expression enhances transcription. Importantly, by focusing on a representative RBP, we delineate the biochemical mechanism by which PSPC1 promotes Pol II engagement and activity in sequential steps. PSPC1 not only prevents the RNA-induced eviction of unphosphorylated CTD, and also synergizes with RNA to promote CTD incorporation, and subsequent phosphorylation and release by CDKs. In addition, PSPC1 stabilizes the binding of the Pol II holoenzyme to template during *in vitro* transcription. Accordingly, auxin-induced degradation of PSPC1 leads to global downregulation of Pol II occupancy and nascent transcription in ESCs. The rescue of defective Pol II binding was not observed in PSPC1 mutants that lack either the major LCS or RRM domain. These multiple lines of evidence corroborate a direct and functional involvement of PSPC1 in transcription through its phase-separation and RNA-binding activities. These two intrinsic properties, which are shared by many chrRBPs, endow PSPC1 with the ability to modulate Pol II binding and transcription activity through its chromatin association.

Based on these findings, we propose that RBPs stabilize Pol II engagement to the transcription sites via RNA and phase separation (Fig. 6). We extrapolate that in cells, the basal activity of Pol II produces short RNA transcripts, which evicts Pol II from gene promoters before the CTD is properly phosphorylated. In the meanwhile, nascent RNA and/or the transcription machineries recruit RBPs to the proximity of the transcription sites via weak and less-specific interactions. These RBPs not only balance negatively charged RNA to protect against the precocious dissolution of Pol II condensates, and also make use of RNA as a multivalent molecule to enhance Pol II phase behavior. During continuous rounds of Pol II fall-off and rebinding, the accumulation of nascent RNA recruits more RBPs until the eventual formation of phase-separated condensates. These RBP-rich condensates concentrate Pol II and necessary enzymes in place to enhance Pol II binding to the transcription sites. Once formed, Pol II is hyper-phosphorylated by CDKs and then released for effective elongation. In this regard, the recruitment of RBPs to chromatin critically contributes to the rate-limiting step of polymerase condensate formation. This model agrees with the observation of dynamic and transient assembly of polymerase clusters in cells^22,25,26,32^.

**Fig. 6.**
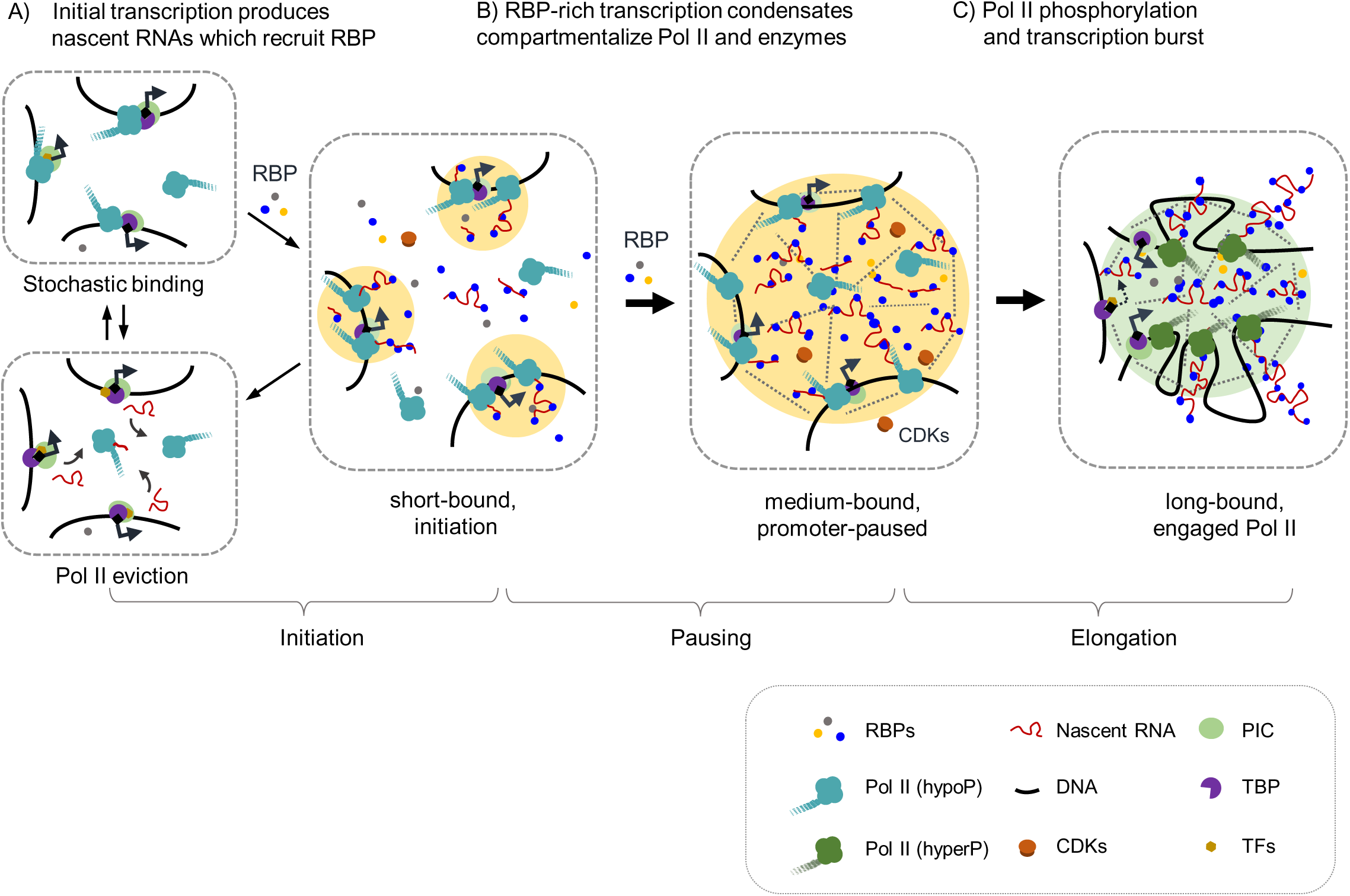
A model depicts that RBPs harness RNA-binding and phase-separation activities in promoting Pol II transcription. Stochastic binding and assembly of Pol II initiates a basal level of transcription. Nascent RNA recruits RBPs to the vicinity of transcription sites. These RBPs not only prevent the premature release of Pol II by RNA (**A**, left), and also utilize multivalent RNA interactions to promote phase separation (**A**, right). When RBPs are scarce or not readily to be recruited, extra repulsive-like charges on RNA dissolve the condensates, leading to Pol II release from the promoter. Iterative cycles of Pol II fall-off and re-binding may occur during the initiation and promoter-proximal pausing until a sufficient number of RBPs are recruited. At the transcription sites, increased molecular crowding further enhances the phase separation to concentrate Pol II and necessary enzymes such as CDKs (**B)**. The RNA-binding activity and multivalent interactions within the transcription condensates stabilize the association of RBPs with transcription sites, and also empower RBPs to stabilize Pol II engagement during initiation and pausing. Formation of large RBP-rich transcription condensates eventually facilitates Pol II phosphorylation by CDKs and elongation into the gene body **(C)**.

The abundance and ability of chromatin-associated RBPs to polymerize and bind RNA favor RBPs as the major components that drive the phase separation of transcription condensates under physiological conditions in the nucleus. In addition, co-transcriptional RNA processing deploys multifunctional RBPs to reside in the proximity of transcription sites, which also offers a convenient means for their moonlighting during the assemblage of transcription condensates. By sensing levels of nascent transcripts, RBPs may leverage transcription output to balance cellular activities. Some RBPs, like PSPC1, directly contribute to formation of the transcription condensates via their intrinsic capability to polymerize, while others, like WDR43, modulate the activity of associated enzymes^61^, and yet others merely increase molecular crowding. Nevertheless, these RBPs are actively recruited and play collaborative roles in both forming and running the transcription ‘factories’^27,76^. We propose that the RBP-RNA interplay represents an important layer of gene regulation, expanding the horizon in our understanding of the intricate regulation of transcription and expression heterogeneity in multicellular organisms.

**Supplementary Information** is available for this paper.

## Acknowledgments

We thank the Shen Laboratory members for insightful discussion. This work was supported in part by the National Basic Research Program of China (2018YFA0107604, 2017YFA0504204 to X.S.) and National Key R&D Program of China (2018YFC1004500 to B.L.); the National Natural Science Foundation of China (31925015, 31829003 to X.S.; 32030019, 31872817 to B.L.); and Beijing Advanced Innovation Center for Structural Biology at Tsinghua University (to X.S.).

## Author contributions

X.S. supervised the study. X.S. and W.S. conceived of and designed the experiments. W.S. performed most experiments with the help of X.B., B.G., Z.L., and W.R., and conducted bioinformatics analysis with assistance from Y.X., X.B., and J.L.. X.B. performed ChIP-seq of ^FB(EXO)^PSPC1, UTP3, UTP6, and CIRH1A. *In vitro* transcription system was designed and set up by B.L. and J.W.. W.S. performed *in vitro* transcription assays with the help of Y.P.. Y.Y. performed ChIP-seq of SAFB1, SAFB2 and hnRNPU. W.Z., X.J., and H.D. provided technical assistance/suggestions for mass spec analysis. Z.W., K.W., G.Z., T.L., and J.W. contributed assistance/suggestions for experiments. X.H. performed ESC total proteome analysis. X.S. and W.S. wrote the manuscript with input from all authors.

## Author Information

The authors declare no competing financial interests. Sequencing data have been deposited in the GEO database under the accession number GEO: GSE150399.

## Methods

### Experimental model and subject details

Mouse ESCs (CJ9, 46C lines and cells expressing endogenous or exogenous 3 × FLAG- and biotin-tagged RBPs) were cultured in complete ESC medium, which includes DMEM (Dulbecco’s modified Eagle’s medium) supplemented with 15% heat-inactivated FCS (fetal calf serum), Penicillin-Streptomycin Solution (100× stock, Life Technologies), 2 mM Glutamax (100× stock, Life Technology), 1% nucleoside mix (100× stock, Millipore), 0.1 mM non-essential amino acids (Gibco), 0.1 mM 2-mercaptoethanol (Gibco) and supplied with 1000 U/ml recombinant leukemia inhibitory factor (LIF, Millipore). ESCs were cultured on plates which were pre-coated with 0.1% gelatin. ESCs used in this study are male. The HEK 293T cells were cultured in medium containing DMEM, 10% FCS, 1× Penicillin-Streptomycin Solution.

### Cells and Culture

Mouse ESCs, including the wild-type (CJ9, 46C lines) and cells expressing BirA with either endogenous or exogenous 3 × FLAG-biotin-tagged proteins, were maintained in complete ESC culture medium: DMEM (Dulbecco’s modified Eagle’s medium) supplemented with 15% heat-inactivated FCS (fetal calf serum), 2 mM Glutamax (100× stock, Life Technology), Penicillin-Streptomycin Solution (100× stock, Life Technologies), 0.1 mM nonessential amino acid (Gibco), 1% nucleoside mix (100× stock, Millipore), 0.1 mM 2-mercaptoethanol (Gibco) and supplied with 1000 U/ml recombinant leukemia inhibitory factor (LIF, Millipore). 293T cells were cultured in DMEM supplemented with 10% FCS.

### Construction of ^AID-FB(KI)^*PSPC1* ESCs

The usage of the auxin-induced degron (AID) system was based on a previous report^76^. ^AID-FB(KI)^*PSPC1* ESCs were constructed by knocking a 3× FLAG-biotin-AID-tag into the 5’ end of the *PSPC1* gene locus. Specifically, we co-transfected wild-type ESCs (CJ9) with one sgRNA targeting the first exon of *PSPC1* (puromycin-resistant) and vectors expressing CRISPR/CAS9 with the plasmid harboring the 3 × FLAG-biotin-AID tag flanked by two homologous arms surrounding the knock-in sites (neomycin-resistant). After selection with puromycin and neomycin for 3 days, single colonies were picked and positive colonies were identified by PCR genotyping.

Before IAA treatment, we infected cells with fresh TIR1 virus, which encodes a protein that mediates ubiquitination and degradation of AID-tagged proteins^76^. Cells were treated with blasticidin (the antibiotic resistant gene carried by TIR1 vectors) for 5 days to select for efficient depletion. We noticed that cells with integrated TIR1 became adapted to IAA after repeated passages, so we used knock- in ESCs at early passage, and we freshly infected them with TIR1 virus right before experiments involving IAA treatments. The final concentration of IAA (Sigma, I5148) in all assays was 1 mM.

### Construction of cells with stably expressed FLAG-biotin-tagged RBPs

There is limited availability of antibodies suitable for co-IP and ChIP-seq analyses; therefore, we constructed ESCs that stably express FLAG-biotin-tagged RBPs as previously described^77^. The cDNA of RBPs (PSPC1, hnRNPU, SAFB, SAFB2, UTP3, UTP6 and CIRH1A) fused with 3 × FLAG-biotin tag cDNA was cloned into PiggyBac vectors. The PiggyBac vectors expressing these proteins were co-transfected with pBase vector into J1 ESCs expressing the bacterial biotin ligase birA, and stable clones were selected by treatment with hygromycin.

### Chromatin fractionation and mass spectrometry analysis

Four 15-cm plates of ESCs were harvested and washed with cold PBS. Five pellet volumes (PVs) of hypotonic buffer (20 mM HEPES, pH 7.5, 10 mM KCl, 1.5 mM MgCl_2_, 1 mM EDTA, 0.1 mM Na_3_VO_4_, 0.1% NP-40) supplemented with proteinase inhibitors were added to cell pellets, which were then transferred to a pre-chilled 15 ml Dounce tissue homogenizer (Wheaton Scientific). The cells were gently homogenized up and down 10 times and then spun for 5 minutes at 1,300 g, 4 °C. The pelleted nuclei were subjected to crosslinking by 1% formaldehyde for 10 minutes and the reaction was ended by adding 1/20 volume of 2.5 M glycine. Next, the crosslinked nuclei were resuspended with 2 × pellet volumes of nuclear lysis buffer (50 mM Tris-HCl, pH 8.1, 10 mM EDTA, 1% SDS) and incubated on ice for 10 minutes. 0.5 volume of ethanol was added and the DNA-protein complexes were precipitated at −20 °C for 1 hour. The DNA-protein complexes were spun down at 5,000 g at 4 °C for 20 minutes. The pellet was further washed with ice-cold 75% ethanol and resuspended in 50 mM Tris-HCl buffer (pH 7.4). Urea (final 8M) and SDS (final 2%) were added to the suspension, and the mixture was incubated at 37 °C for 30 minutes with gentle shaking. An equal volume of 5 M NaCl was added and the resulting mixture was incubated at 37 °C for another 30 minutes. The DNA and its associated proteins were precipitated again by the addition of 0.1 volume of 3 M sodium acetate and 3 volumes of ice-cold ethanol. Precipitated DNA and DNA-protein complexes were collected by centrifugation at 5,000 g at 4 °C for 5 min and washed twice with ice-cold 75% ethanol to remove salts and detergents. The pellet was air-dried and resuspended in DNase digestion buffer (20 mM HEPES, pH 7.5, 15 mM NaCl, 6 mM MgCl_2_, 1 mM CaCl_2_, 10% glycerol) containing DNase I (10 U, Takara) and incubated at 37 °C for 1 hour. EDTA was added to end the reaction and the pellet was spun down at 13,000 rpm for 20 minutes at 4 °C. Proteins released into the supernatant by DNase treatment were collected and subjected to SDS-PAGE. Proteins migrating above 20 kD (to exclude histones) were collected for mass spec sequencing.

Raw peptide information was used for protein identification by the MaxQuant platform and the protein abundance was valued by the iBaq intensity^78^. We took the proteins identified by two replicates with both iBaq intensity > 500 and molecular weight > 20 kD as chromatin proteins. In order to measure the relative abundance of different proteins and compare between different batches of experiments, we defined an iBaq ratio by normalizing each protein’s iBaq intensity to the sum of all proteins’ iBaq intensities. Gene classification was based on gene ontology analysis (GO). Proteins that are involved in multiple biological processes were marked with the biological term that ranks higher in GO analysis.

### Salt extraction of native chromatin

Native nuclei were isolated as described above in hypotonic buffer and divided equally into 4 tubes. Four volumes of extraction buffer (20 mM HEPES pH 7.5, 10 mM KCl, 1.5 mM MgCl_2_, 1 mM EDTA, 0.1 mM Na_3_VO_4_, 25% glycerol, 1 mM PMSF, 1/200 Proteinase Inhibitor cocktail) with different concentrations of NaCl from 200 to 500 mM, were added separately to each tube and rotated at 4 °C for 30 minutes. The nuclei were then spun at 14,000 rpm for 20 min at 4 °C. The supernatant represents the extracted nuclear fraction and the pellet represents the native chromatin resistant to salt extraction. The same percentage (5%) of supernatant and pellet was used for western-blot. The antibodies used are listed here: WDR43 (Abclonal, Q659), hnRNPU (Abcam, ab180952), hnRNPL (Santa Cruz, sc-32317), PTBP1 (Abclonal, A6107), DDX5 (Abcam, ab126730), FUS (Abcam, ab70381), PSPC1 (Abcam, ab104238), pan-Pol II (Abcam, ab52202), EZH2 (CST, 5246S), SUZ12 (CST, 3737S), CTCF (Abcam, ab128873), CHD1 (CST, 4351S), TOP2A (Abcam, ab52934), and H3 (Easybio, BE3015).

### Quantitative mass-spectrometry (MS) after transcription inhibition or RNase treatment

Cells were treated separately with actinomycin D (ActD, 1 μg/ml for inhibition of both Pol I and Pol II transcription, 10 ng/ml for inhibiting only Pol I transcription, Abcam, ab141058), triptolide (TPL, 1 μM, Abcam, ab120720) or DMSO for 2 hours before chromatin fractionation. RNase A treatment was performed as previously described^79^. Briefly we used PBS with 0.05 % triton to permeabilize cells at room temperature for 2 minutes. Cells were then quickly spun at 1,200 rpm for 3 minutes at 4 °C. Permeabilized cells were then mock treated or treated with 1 mg/ml RNase A (Takara) diluted in PBS for 20 minutes at room temperature. Cells were spun down and washed with PBS for later fractionation.

For the Label-free Quantification (LBQ) method, we performed MS analysis under 4 experimental conditions (DMSO vs ActD, Mock vs RNase) with one replicate for each and we quantified each protein’s relative abundance by iBaq intensity as described above. For the Tandem Mass Tag (TMT) method, there were 5 experimental conditions analyzed (DMSO vs ActD or TPL, Mock vs RNase) with one replicate for each. We performed the experiment as previously published^80^. After chromatin fractionation, we used the same amount of chromatin proteins for different conditions and labeled them with different amine-reactive TMT 6-plex reagents (ThermoFisher). Then we mixed these samples together and carried out mass spectrometry analysis. Lastly, for Stable Isotope Labeling with Amino Acids (SILAC), cells cultured with heavy SILAC media were firstly treated with transcription inhibitor or RNase and mixed with an equal number of mock cells cultured in light media. We exchanged the media for different treatments for an additional biological replicate to exclude media bias. Mixed cells were used for chromatin fractionation and mass spec analysis.

The majority (> 91%) of the defined set of 512 chrRBPs were identified by the LBQ method. For TMT and SILAC, only 50% to 90% of chrRBP hits were detected with quantitative information among 9 samples. It is possible that not all of the 512 chrRBPs were detected by TMT and SILAC because these two MS methods involve isotope labeling. In addition, we noted that transcription inhibition or RNase treatment resulted in decreased abundance of some proteins. Thus, we reason that combined effects of labeling efficiency and decreased protein abundance may contribute to the limited protein detection by TMT and SILAC. Nevertheless, all the 512 chrRBPs were identified using at least one method.

In order to enable cross-comparison between experiments with different MS methodology and quantification, we first calculated fold-change (FC) scores by normalizing the experimental (exp) sample to the corresponding mock treatment as below. LBQ: log5 (exp / mock + 0.001); TMT: log3 (exp / mock + 0.001); SILAC: log2 (exp / mock + 0.001). For each treatment, we then calculated the mean of normalized FC scores by three methods and used a cutoff lower than −0.2 to select chrRBPs that are dynamically regulated by transcription/RNA.

The validation of quantitative MS was performed as described above. To distinguish effects of Pol I transcription from Pol II transcription, we added an extra group with low concentration of ActD (10 ng/ml) treatment for 2 hours that only inhibits Pol I transcription. Chromatin fraction and total lysates were collected from same samples. Additional antibodies used for western-blots are listed here: DDX21 (Novus Biologicals, NBP1-83310), FUBP1 (Abcam, ab181111), FUBP3 (Abcam, ab181025), LIN28A (Abcam, ab155542), NCL (Abcam, ab134164) and TUBULIN (CWBIO, CW0098).

### Analysis of biochemical features of chrRBPs versus non-chrRBPs

We analyzed the biochemical features of chrRBPs by using previously developed methods or available website tools.

Analysis of low-complexity sequence (LCSs): http://repeat.biol.ucy.ac.cy/fgb2/gbrowse/swissprot/81; Analysis of intrinsically disordered regions (IDR): http://www.pondr.com/82, https://iupred2a.elte.hu/83, https://github.com/zhanzhan90/distribution-of-amino-acid.git (DOI:10.5281/zenodo.3874019); Analysis of isoelectric point (pI): https://web.expasy.org/compute_pi/; RNA-binding domain (RBD)^84^.

### Co-immunoprecipitation (co-IP) and biotin-mediated affinity purification (bio-AP)

One 10 cm plate of ESCs was harvested and washed twice with cold PBS, and lysed in 5 volumes of IP lysis buffer (50 mM Tris pH 7.4, 150 mM NaCl, 0.5% TritonX-100, 10% glycerol, 1 mM DTT, 1 mM PMSF and 1/200 Proteinase inhibitor cocktail) supplemented with 1 μl benzonase (Sigma) at 4 °C with rotation for 30 minutes. The lysate was later cleared by centrifugation for 20 minutes at 14,000 rpm, 4 °C. 5% of the lysate was collected as input. For antibody IP, 2-3 μg antibody/IgG were added to the cleared lysate and incubated overnight. 25 μl pre-equilibrated ProteinA/G resins (ThermoFisher 53133) were added and incubated for another 3 hrs. For biotin-mediated affinity purification, 30 μl pre-equilibrated M-280 dynabeads (Invitrogen) were added instead and incubated at 4 °C overnight. The beads were washed with IP lysis buffer three times and eluted with SDS loading buffer. One third of the sample was loaded for western-blot analysis. Additional antibodies used are listed here: SNRNP70 (Santa Cruz, sc-390988), hypo-phosphorylated Pol II (8WG16) (Covance, MMS-126R), Pol II Ser5P (CST, 13523), Pol II Ser2P (CST, 13499), TBP (Santa Cruz, sc-421) and FLAG (Sigma-Aldrich, F3165).

### Immunofluorescence (IF)

Cells grown on matrigel-treated coverslips were fixed by 4% paraformaldehyde (PFA) for 15 min at room temperature, followed by blocking and permeabilization with blocking buffer (PBS supplemented with 5% BSA and 0.5% TritonX-100) for 45 minutes at room temperature. Antibodies were diluted in blocking buffer and incubated for 1 hour at room temperature. Dilution was based on the manufacturer’s instruction: PSPC1 (Abcam, ab104238, 1:100), pan-Pol II (Abcam, ab52202, 1:100), TBP (Santa Cruz Biotechnology, sc-421, 1:100). After washing three times with PBS for 5 min each, fluorescent secondary antibody (1:1000) diluted in blocking buffer was added and incubated for 45 minutes at room temperature. The cells were mounted in Fluoromount-G (SouthernBiotech). Pictures were taken with a Nikon A1R-HD-Multiphoton microscope.

### Fluorescence recovery after photobleaching (FRAP)

Cells were transfected with mCherry-fused PSPC1 or GFP-fused H3. After 24 hours post-transfection, the cells were plated on matrigel-treated glass-bottom confocal Petri dishes (CELLVIS). A Nikon A1R-HD-Multiphoton microscope was used for photobleaching, and quantitation was performed as previously described^85^.

### Protein purification in bacteria

Protein purification in bacteria was performed as described^77^. Briefly, the expression plasmids for His-tagged PSPC1, mCherry-PSPC1, mCherry-PSPC1_ΔLCS2_, mCherry-PSPC1_RRMmut_, GFP-CTD, and SNAP-TBP were transformed separately into *E. coli* (DE3). Cells were cultured in 6 L of LB media at 37°C until OD_600_ reached 0.6. Protein expression was induced by addition of 0.5 mM IPTG, and cells were cultured at 18°C overnight. Cells were lysed in a buffer containing 500 mM NaCl, 50 mM Tris (pH 8.0), 2 mg/mL lysozyme, then sonicated and centrifuged at 18,000 rpm at 4°C for 1 hour. The supernatant was incubated with Ni-NTA resin and washed with a buffer containing 1 M NaCl, 20 mM Tris (pH 8.0) and 20 mM imidazole. Protein was eluted in a buffer containing 500 mM NaCl, 50 mM Tris (pH 8.0) and 250 mM imidazole. The protein solution was diluted with lysis buffer, followed by Source Q/S, or Heparin or/and Superdex 200 column. Purified proteins were concentrated in concentration tubes with the buffer (20 mM Tris pH 8.0, 150-500 mM NaCl) and flash-frozen in liquid nitrogen and stored at −80°C.

### Purification of CDK7 and CDK9 complexes

For the purification of CDK7, 293F cells were cultured in SMM 293-TI serum-free medium (Sino-Biological, M293TI). The cells were split to a density of ∼ 10^6^ cells/ml 24 hours before transfection, then further cultured with shaking at 37℃. ∼200 ml 293F cells were used for transfection. 200 µg pMlink-StrepII-FLAG-CDK7 plasmids were diluted with 10 ml DMEM medium. 600 µg Polyethylenimine (PEI) were diluted with another 10 ml DMEM medium. The plasmid solution and PEI solution were gently mixed together and incubated for 30 minutes at room temperature. Then the mixed solutions were gently added into the 293F cell cultures. The transfected cells were further cultured for 48 hours and harvested by centrifugation. The cells were washed twice with PBS and then lysed in 20 ml lysis buffer (50 mM Tris pH 7.4, 150 mM NaCl, 0.5% Triton X-100 and 10% glycerol) supplemented with 1 mM PMSF and 1/500 Protease inhibitor cocktail at 4 ℃ for 30 minutes. Insoluble fractions were removed by centrifugation at 12,000 g for 20 minutes at 4℃. The lysate was incubated with 500 µL Strep Tactin beads (pre-equilibrated with lysis buffer) for 1 hour. The beads were washed 5 times with high salt wash buffer (50 mM Tris pH 7.4, 350 mM NaCl, 0.5% Triton X-100 and 5% glycerol). The proteins were eluted by 2.5 mM desthiobiotin (dissolved in 50 mM Tris pH 7.4, 150 mM NaCl, 10% glycerol). The eluted proteins were concentrated by a Millipore concentration tube (< 10 KD).

6× His-tagged human CDK9 and CyclinT1 baculoviruses were gifts from Guohong Li’s lab. The baculoviruses were amplified and infected into cells as described by the manual in the Bac-to-Bac Baculovirus Expression System (Invitrogen). ∼200 ml Sf9 cells were infected for 60 hours. The cells were lysed by 20 ml lysis buffer (50 mM Tris pH 7.4, 150 mM NaCl, 0.5% Triton X-100, 10% glycerol and 10 mM imidazole) supplemented with 1 mM PMSF and 1/500 Protease inhibitor cocktail. The lysate was centrifuged at 12,000 g for 20 minutes at 4℃. The supernatant was further incubated with a Ni^2+^ beads column. The column was washed with 30 ml wash buffer (50 mM Tris pH 7.4, 350 mM NaCl, 0.5% Triton X-100, 5% glycerol and 20 mM imidazole). The proteins were eluted by 300 mM imidazole buffer (50 mM Tris pH 7.4, 150 mM NaCl, 10% glycerol). Imidazole was removed by buffer exchange during protein concentration with a Millipore concentration tube. Proteins were stored in storage buffer containing 50 mM Tris pH 7.4, 150 mM NaCl and 10% glycerol.

### Estimation of protein molecule number and nuclear concentration

The nuclear concentrations of proteins were calculated as previously described in HeLa cells^86,87^. Briefly, the nuclear concentration of a nuclear protein equals the protein molecule number divided by the nucleus volume ^87^. The nuclei volume of a HeLa cell was assumed to be 220 fl as previously measured ^86,88^. We presume that chrRBPs are mainly localized in nuclei and their nuclear concentrations may be in similar ranges between Hela cells and ESCs. In Hela cells, the estimated nuclear concentrations are 0.65 μM for Pol II, 3.3 μM for PSPC1, and 0.06 μM for TBP. In addition, we also referred the published work in ESCs ^89^ and estimated the nuclear concentration of TBP protein to be ∼0.3 μM by assuming that an ESC nucleus is 10 µm in diameter. The molecule number and nuclear concentration of each protein is listed in Supplementary Table 1.

### *In vitro* droplet formation

*In vitro* droplet formation was performed in tubes and visualized in 384-well glass-bottom plates (In Vitro Scientific, P384-1.5H-N). Proteins were diluted to indicated concentrations in a buffer containing 20 mM Tris pH 7.5 and 150 mM NaCl supplemented with or without 10% dextran.

For imaging, we mixed mCherry-PSPC1 or Cy5.5 NHS ester labeled PSPC1 with untagged PSPC1 at the ratio of 1:3 or 1:10 as the concentration of purified mCherry-PSPC1 is relatively low. SNAP-TBP was firstly labeled with SNAP-Surface® Alexa Fluor® 647 following the manufacturer’s instructions and then mixed with unlabeled protein (1:10) for imaging. For phase-separation assays with RNA, all buffers used were kept RNase-free and supplemented with RNase inhibitor. Total RNA was extracted from ESCs and added as indicated. All pictures were taken with a Nikon confocal microscope at the same time and analyzed by ImageJ. Total fluorescence intensity of CTD was obtained by calculating the sum of CTD fluorescence intensity in droplets for each field of view. We took 10 pictures in different views for each condition, then used the images for statistical analysis.

For droplet sedimentation, samples were centrifuged for 10 min at 14,000 rpm, 4 °C. The same fraction of supernatant and pellet was used for western-blot analysis. Anti-GFP antibody (Santa Cruz, sc-9996) was used for detecting GFP-fused CTD.

### Kinase assays

Kinase assays were performed as previously described ^90^ with modifications. About 0.2 μg of GFP-CTD was pre-incubated with the same amount (1-5 μg) of mCherry-PSPC1, or mCherry-PSPC1_ΔLCS2_, or mCherry-PSPC1_RRMmut_ for 20 minutes at room temperature in kinase buffer (20 mM Tris-HCl pH 7.0, 150 mM KCl, 2 mM MgCl_2_, 2 mM DTT, 0.15 mg/mL BSA). mCherry or BSA were used as controls. For kinase assays with RNA, all buffers were kept RNase-free and supplemented with RNase inhibitor. Total RNA from mouse ESCs was added to a final concentration of 50 ng/μl. Then, 0.2 μg of CDK9 or CDK7 and ATP (final 0.1 mM) were added and incubated at room temperature for 10 minutes. SDS loading buffer was added to end the reactions. Pol II Ser5P antibody (CST, 13523) was used for detecting phos-CTD, and mCherry antibody (CST, 43590) was used for detecting mCherry-fused PSPC1 mutants.

### CTD release assay

TBP (5 μM) and CTD (0.6 μM) were firstly mixed together with PSPC1 (5 μM) or mCherry (5 μM) in a buffer containing 20 mM Tris pH 7.5, 150 mM NaCl, and 10% dextran before imaging or sedimentation experiments.

For imaging, upon addition of ATP (0.1 mM) and CDK9 (2 μg), the plate was immediately put under the microscope and recording was started. For every sample, at least 2 views were recorded at the same time. For assays with RNA, we noticed that addition of RNA greatly accelerated the release of CTD. To capture the release of CTD and slow down the reactions, we used a less amount of CDK9 (1 μg). Time-lapse analysis of droplet intensity analysis was performed using Nikon NIS-element AR software. Briefly, we selected droplets that were already present at the start and quantified the mean intensity of the selected droplets at each time point. To avoid environmental disturbances, we normalized each droplet’s CTD mean fluorescence intensity to its TBP mean fluorescence intensity. To compare across different samples, we further normalized to the initial levels at the start, and obtained the normalized CTD intensity curve i (t) for each droplet. To calculate CTD release rate, we did non-linear fitting of the intensity curve using GraphPad Prism. To simplify the mathematical calculation, we assumed that the droplet was a homogenous sphere. For each droplet, we drew an intensity curve i(t) and then derived the CTD release rate based on two equations: v (t) = ΔI (t) / (4πR^2^); I (t) = 4πR^3^ / 3 × i (t). Therein, v, t, I, i, ΔI and R, respectively, represent the release rate, time, total fluorescence intensity, mean fluorescence intensity, the change of total fluorescence intensity and droplet radius. Thus, we can get the rate curve v (t) = R / 3 × Δi (t).

For droplet sedimentation, we fractionated droplets at each indicated time. The same fraction of the supernatant and pellet was used for western-blot.

### Preparation of Bubble-601R DNA (BR) template

Bubble DNA was produced by annealing either unlabeled forward primer (5’[Cy5.5]-AGGCAGGCCTTAGCTCCGTTCGCCGTGTCCTACCTATCCTCTCCTCACCACTCCCGGGGCC ATTC) or an 5’Cy5.5 labeled forward primer of the same sequence with one 5’ phosphorylated reverse primer (5’[phos]TGGCCCCGGGAGTGGTGAGGAGAGGATAGGTAATCAGTTACGCCCGGAGCTAAGGCCTGCCTAGT). The resulting fragments bear a Bgl I site at the downstream end (Supplementary Fig. 5a). 601-R fragments were generated through Bgl I and Dra III (NEB) double digestion of pJW013, which contains 10 copies of the 601-positioning sequence and a linker region and purified using Model 491 Prep Cell (Bio-Rad). Annealed bubble templates and digested 601-R fragments were ligated using T4 DNA ligase (NEB) and ligated products were further purified through Model 491 Prep Cell to remove free bubbles and 601R fragments.

Plasmids used in this assay

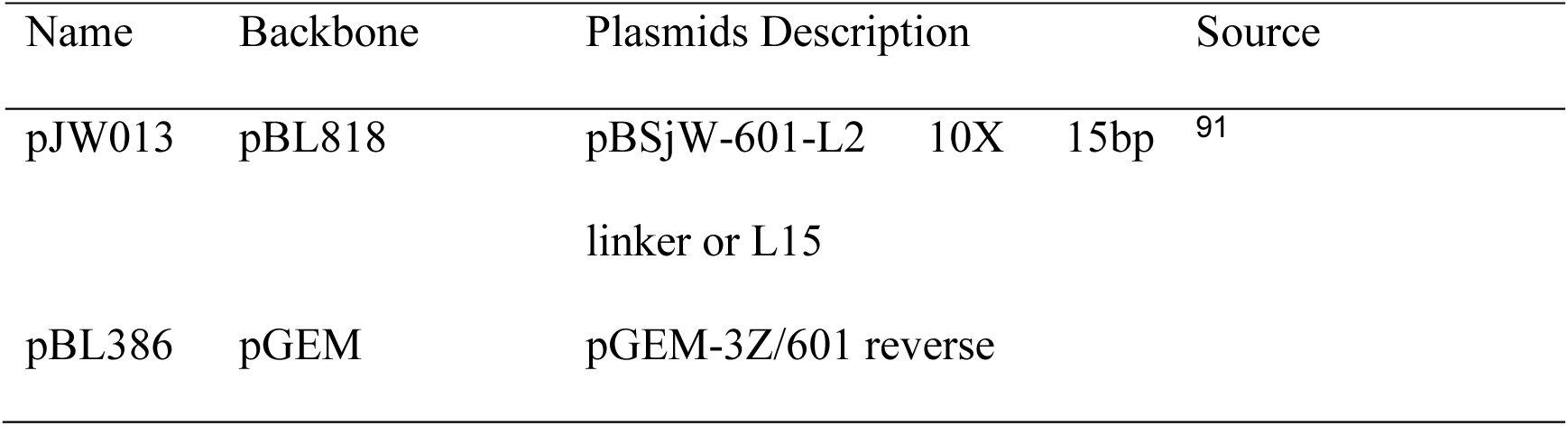

### Purification of Pol II

Pol II complexes were purified through tandem affinity purification (Rpb9-TAP) as described ^92,93^ with minor modification. Briefly, 6 liters of yeast culture (YBL360) were grown in YPD medium at 30°C. Cell pellet was resuspended in an extraction buffer (AE buffer) (40 mM HEPES.KOH pH7.5, 350 mM KOAc, 10% glycerol and 0.1% tween 20, supplemented with complete sets of fresh proteinase inhibitors) and lysed using a bead beater (Biospec). Homogenized cell suspension was then treated with 75 μL of 10 mg/mL heparin (Sigma) and 75 μL of DNase I (Sigma) to facilitate releasing Pol II from genomic DNA. Resulting extracts were clarified through ultracentrifugation and directly applied to standard TAP purification ^93^ using IgG Sepharose and calmodulin resin (GE). Pol II complexes were eluted with Acetate Calmodulin Elution Buffer (ACEB) (10 mM Tris.OAc pH8.0, 150 mM KOAc, 1 mM Magnesium acetate, 1 mM imidazole, 2 mM EGTA pH8.0, 10 mM BME, 0.1% NP40 and 10% glycerol).

Yeast strain used in this assay:

| Name | Parental strain | Genotype | Source |
| --- | --- | --- | --- |
| YBL360 | BY4741 | MATa his3Δ1 leu2Δ0 met15Δ0 ura3Δ0Rpb9-TAP | OpenBio<sup>92</sup> |

### *In vitro* transcription and gel-shift assays

*In vitro* transcription reactions were carried out in 20 μL transcription buffer (Buffer B), which contains 25 mM HEPES pH7.5, 50 mM KCl, 10% Glycerol, 5 mM MgCl2 (Sigma), 1 mM DTT and 0.05 mg/ml BSA (Sigma). To detect transcribed RNA, 0.4µl of Low-C NTP mix (25 mM ATP, 25 mM UTP, 25 mM GTP and 0.25 mM CTP) and 0.5 µl of α-32P-CTP (3000 Ci/mmol, 10 mCi/ml Perkin Elmer) were supplemented into each reaction along with 1 ng of DNA template and 100 ng of competitor DNA. Reactions were incubated at 30°C for 45 mins and then stopped by adding 120µl of STOP buffer (0.3 M NaAC, 5 mM EDTA, 0.1% SDS, 40 µg/ml linear acrylamide (Ambion 9520)) and 1 µl of Proteinase K (20mg/ml). The mixtures were placed at 55°C for 15 mins and then subjected to phenol/chloroform extraction and ethanol precipitation. Pellets were resuspended using 12 µl of 90% Formamide-TBE Loading Buffer and denatured before being loaded onto a 8% polyacrylamide gel (19:1) containing 7 M Urea. Dried gel was exposed to Phospho-imager and scanned with GE Typhoon Scanner. For gel-shift assays, 50 ng of Cy5.5-labeled bubble-601R template and 25 mM NTPs mix were used in otherwise similar transcription condition described above. For the heparin treatment, 2.3 μL of 250 ng/μL heparin or mock was added to each reaction and incubated at 4 °C for 20 min. Samples were directly loaded onto a 3.5% native polyacrylamide gel (37.5:1) in 0.3 x TBE. Electrophoresis was carried out at 4°C for 3.5 hours and gels were scanner with Li-Cor CLX scanner.

### ChIP-qPCR and ChIP-seq analysis

ChIP assays for endogenous proteins, including Pol II Ser2P (CST, 13499) and Pol II Ser5P (CST, 13523), were performed as described ^94^. For RBPs, we performed FLAG or biotin-mediated ChIP-seq or ChIP-qPCR with endogenously or exogenously expressed FLAG-biotin-tagged proteins, due to the lack of ChIP-grade antibodies. Cells were subjected to single-step or tandem ChIP analysis under standard or strong crosslinking conditions. For ^FB(OE)^PSPC1, cells were crosslinked by 1% FMA for 10 minutes. For ^AID-FB(KI)^PSPC1, as well as exogenously expressed UTP3, UTP6 and CIRH1A, harvested cells were crosslinked by 3% formaldehyde for 10 min. For each ChIP-seq, 5 μg FLAG antibodies (Sigma F1804) were used for each experiment and 25 μl slurry Protein A/G UltraLink Resins (ThermoFisher 53133) were used for each IP. Specifically, for hnRNPU, SAFB and SAFB2, cells were crosslinked by 2 mM DSP (dithiobis succinimidyl propionate) for 30 minutes, followed by a 10-minute 1% formaldehyde crosslinking. The crosslinked cells were first partially fragmented by 12 U/ml DNase I at 37 °C for 10 minutes, then sonicated at 25% amplitude for 30 seconds. After FLAG antibody IP, these samples were subjected to a second purification step using 30 μl M-280 Streptavidin Dynabeads (Invitrogen 11205D). The rest of the steps were performed as previously described^94^. The ChIP-seq library was constructed using an NEBNext® ChIP-Seq Library Prep Reagent Set or using Tn5 following ChIPmentation protocol ^95^, and sequenced on an Illumina Hiseq 2500 or X10 platform.

ChIP-seq analysis was performed as previously described^77^. The reads were aligned to the mouse genome (NCBI build 37, mm9) using the bowtie2 program with default parameters. Peaks were called using the MACS program (*p* < 10e-3). Annotation of the peaks was completed using the ‘annotatePeaks’ module in the HOMER program. ChIP-seq peaks located within 5 kb around transcription start sites were defined as promoter peaks. For RBP co-occupancy analysis, 14 analyzed chrRBPs co-occupy at a total of 15,317 promoters, which belong to 75% of annotated protein-coding genes (20,516) in the mouse genome (see Supplementary Table 4). Of note, proteins that were analyzed in ectopically and endogenously tagged forms, for example PSPC1 (Extended Data Fig. 6a) and WDR43^77^, share similar sets of ChIP-seq targets, thus excluding a potential effect of ectopic expression.

For analysis of enhancers and super-enhancer, typical enhancers and super-enhancers in mouse ESCs were defined previously^96^. For metagene analysis, gene bodies, typical enhancers and super enhancers were split into 100 bins and the flanking 2 kb regions were split into 20 bins. The read number in each bin was counted and normalized to the length. Heatmaps of ChIP-seq read density were visualized using Treeview 3.0. For clustering analysis of RBP ChIP-seq, read counts in a region encompassing 5 kb around the peak center of all sites bound by RBPs were calculated, and used for Pearson correlation analysis between ChIP-seqs. Unsupervised clustering was analyzed with R.

### Nuclear run-on

Nuclear run-on was performed as previously described^97^ with some modifications. Harvested cells were firstly washed with PBS and permeabilized with NP-40 lysis buffer (10 mM Tris-HCl, pH 7.4, 3 mM MgCl_2_, 10 mM NaCl, and 0.05% NP-40) on ice and quickly spun down at 300 g for 4 minutes. The pellet was resuspended with 40 μl nuclei storage buffer (50 mM Tris-HCl, pH 8.3, 5 mM MgCl_2_ and 0.1 mM EDTA, 40% glycerol). The same volume of 2 × transcription buffer (300 mM KCl, 20 mM Tris-HCl, pH 8.3, 5 mM MgCl_2_, and 4 mM DTT, supplemented with 2 mM each of ATP, GTP and CTP, 1 mM UTP and 1 mM BrUTP) was added and incubated at 30 °C for 10 minutes. RNA was then extracted by Trizol. 2 μg of anti-BrdU monoclonal antibody (Santa Cruz, sc-32323) were incubated with 30 μl pre-equilibrated protein G dynabeads (ThermoFisher) in PBST for 10 minutes at room temperature. Conjugated beads were then blocked by blocking buffer at room temperature for 30 minutes and washed twice by PBSTR buffer (PBST supplemented with RNase Inhibitor). Beads were resuspended in 100 μl PBSTR and added to an equal volume of purified RNA. After incubation at room temperature for 30 minutes, beads were washed three times with PBSTR and the bead-bound RNAs were extracted by Trizol. The newly transcribed RNAs were then reverse transcribed and quantified by RT-qPCR.

### Transient transcriptome sequencing of nascent transcripts (TT-seq)

TT-seq was performed as described previously with some modifications^98,99^. Cells were labeled in media for 10 minutes with 500 µM 4-thiouridine (4sU, Sigma-Aldrich, St. Louis, MO, USA.). RNA extraction was performed with TRIzol (Life Technologies, Carlsbad, CA, USA). Labelled RNAs were further biotinylated and fragmented as previously described. The fragmented RNAs were then subjected to two times of affinity purification. After first time affinity capture by M-280 dynabeads (Invitrogen), beads were washed 2 times (5-10 min each) with 0.5 ml of washing buffer (100 mM Tris pH 7.4, 10 mM EDTA, 1 M NaCl, 0.1% Tween-20) at 45 ℃, followed by another 2 times of wash with 0.5 ml of SDS washing buffer (50 mM Tri-Cl, pH8.1, 10 mM EDTA, 1% SDS) at room temperature. RNA was eluted by 50 μl SDS washing buffer at 95 °C for 5 minutes. Repeat once and combine the eluate. 50 μl pre-washed M280 beads were added to the eluate and incubated for another 20 minutes with rotation at room temperature. Beads were washed as described above. RNA was eluted with 100 µl SDS washing buffer supplemented with 0.1 M dithiothreitol (DTT) and purified as reported. RNA-seq libraries were constructed using NEBNext Ultra II directional RNA library prep kits (NEB).

RNA-seq analysis was performed as previously described^99^. The clean reads were mapped to the mouse genome (mm9) through TopHat. Metagene analysis was performed using ngs.plot^100^. To compare among different samples, we calculated the read density by normalizing to the reads that are mapped to rRNA gene *RN45S*.

### EU incorporation using Click-iT technology

Lentivirus-mediated RNAi (pLKO) was performed as previously described^77^ to knock down RBPs. sh*Ctrl* and sh*RBP* lentiviruses were packaged and generated in 293T cells. Cells were infected three times with lentivirus to achieve efficient knockdown. Infected cells were selected by puromycin for 36 hours at 24 hours post-infection. To label nascent transcripts, cells were labelled for 20 minutes with EU (5-ethynyl uridine, Jena Bioscience CLK-N002, final concentration at 1 mM). Harvested cells were firstly labeled with Zombie Aqua^TM^ (BioLegend) to mark dead cells. Cells were then fixed with 4% formaldehyde for 15 minutes at room temperature and permeabilized for 5 minutes with PBS supplemented with 0.5% TritonX-100. Next, the cells were labeled with Alexa 647 using a Click-iT Cell Reaction Buffer Kit (Life Technologies, C10269) following the manufacturer’s instructions. Labelled cells were subjected to fluorescence-activated cell sorting (FACS) analysis. Quantification of EU intensity was performed using FlowJo software. The intensity of each sample was first compared to the non-labeled blank control. The sh*RBP* samples were then normalized to the control sample transfected with sh*Ctrl*.

### Luciferase Assay

RBP cDNA was cloned and fused with Gal4 DNA binding domain (Gal4DBD) in pcDNA3.1 as previously described^77^. 5× UAS sequence was inserted upstream of the E1b minimal promoter in the psiCHECK-2 vector. Luciferase assays were performed in a 24-well plate. To test the cooperative activity of RBPs in transcription, 100 ng of psiCHECK-2 and 200 ng of each pcDNA3.1-GAL4-RBP vector were co-transfected into HEK293T cells. To exclude dosage effects, we co-transfected pcDNA3.1-GAL4-empty vectors so that every well was treated with a total amount of 1 μg GAL4 expression construct. We performed luciferase assays at 36 hours post-transfection, using a Dual-Luciferase® Reporter Assay System (Promega) following the manufacturer’s instructions. The Renilla luciferase activity was measured and normalized to firefly luciferase activity for comparison across different samples.

### Published datasets used in this study

We have used the following published datasets in our analysis. GSM2988821 WDR43 ChIP-seq, GSM2988831 Pol II 8WG16 ChIP-seq, GSM2988824 Pol II Ser2P ChIP-seq, GSM2988827 Pol II Ser5P ChIP-seq^77^; GSM1941467 pan-Pol II ChIP-seq; GSM3713432 hnRNPK ChIP-seq^101^; GSM3407052 SRSF2 ChIP-seq^102^; GSM1893472 NONO ChIP-seq^103^; GSM1693793 DDX21 ChIP-seq; GSM1915715 LIN28A ChIP-seq^104^; GSM2424700 METTL3 ChIP-seq^105^; GSM560347 MED1 ChIP-seq; GSM1082340 OCT4 ChIP-seq; GSM288356 c-MYC ChIP-seq; GSM1082341 SOX2 ChIP-seq; GSM611197 SIN3A ChIP-seq; GSM1023124 TET2 ChIP-seq; GSM918750 P300 ChIP-seq; GSM480162 SUZ12 ChIP-seq; GSM480161 EZH2 ChIP-seq; GSM769008 H3K4me3 ChIP-seq; GSM1000089 H3K27me3 ChIP-seq; GSM1000099 H3K27ac ChIP-seq; GSM769009 H3K4me1 ChIP-seq; GSM1000109 H3K36me3 ChIP-seq.;

### Quantification and statistical analysis

Statistical analyses were carried out using Excel or R (version 3.3.0). Data are presented as mean ± s.d. For box-plot analysis, outliers are not shown in the figures. The statistical tests used are stated in the relevant figure legends.

## Supplementary information

Supplementary Table 1. Abundance analysis of the ESC chromatin proteome. Related to Fig. 1.

Supplementary Table 2. Comparison of chrRBPs vs non-chrRBPs. Related to Fig. 1.

Supplementary Table 3. Summary of effects of transcription inhibitors and RNA degradation on the chromatin proteome. Related to Fig. 1.

Supplementary Table 4. PSPC1 ChIP-seq targets. Related to Fig. 4.

Supplementary Table 5. Summary of ChIP-seq binding of various RBPs at promoters and enhancers.

Supplementary Table 6. Lists of primers, adaptors and reagents in this study.

Supplementary Video 1. PSPC1 promotes CTD release from TBP droplets. Related to Fig. 2.

Supplementary Video 2. RNA promotes CTD release from TBP-PSPC1 droplets. Related to Fig. 2.

Supplementary Video 3. CTD release from TBP, TBP-PSPC1, and TBP-PSPC1-RNA droplets is ATP-dependent. Related to Fig. 2.

## Supplementary Tables

**Supplementary Table 1. Abundance analysis of the ESC chromatin proteome.**

Abundance analysis and classification of ESC chromatin proteome identified by mass spec.

**Supplementary Table 2. Comparison of chrRBPs vs non-chrRBPs.**

Biochemical features of chrRBPs vs non-chrRBPs.

**Supplementary Table 3. Summary of effects of transcription inhibitors and RNA degradation on the chromatin proteome.**

Quantitative information of changes of chromatin proteins after transcription inhibition or RNA degradation.

**Supplementary Table 4. Summary of PSPC1 ChIP-seq targets.**

Targets identified in each PSPC1 ChIP-seq replicate.

**Supplementary Table 5. Summary of ChIP-seq binding of various RBPs at promoters and enhancers.**

ChIP-seq targets identified for various RBPs.

**Supplementary Table 6. Lists of primers, adaptors and reagents in this study.**

Sequence of primers and sgRNAs used in this study.

## Supplementary videos

**Supplementary Video 1. PSPC1 promotes the release of phosphorylated CTD from TBP droplets.**

Real-time movie of CDK9-induced CTD release from TBP droplets in the presence of mCherry or PSPC1 was taken by confocal microscopy at 1.5-minute intervals for 90 minutes. The first 45 minutes of video is shown here. The scale bar is 20 µm.

**Supplementary Video 2. RNA promotes CTD release from TBP-PSPC1 droplets.**

Real-time movie of CDK9-induced CTD release from TBP-PSPC1 droplets in the presence or absence of RNA was taken by confocal microscopy at 6-minute intervals for 120 minutes. The scale bar is 20 μm.

**Supplementary Video 3. CTD release from TBP, TBP-PSPC1, and TBP-PSPC1-RNA droplets is ATP-dependent.**

Real-time movie of CDK9-induced CTD release from TBP, or TBP-PSPC1 or TBP-PSPC1-RNA droplets in the absence of ATP was taken by confocal microscopy at 6-minute intervals for 120 minutes. The scale bar is 20 μm

---

**Extended Data Fig. 1.**
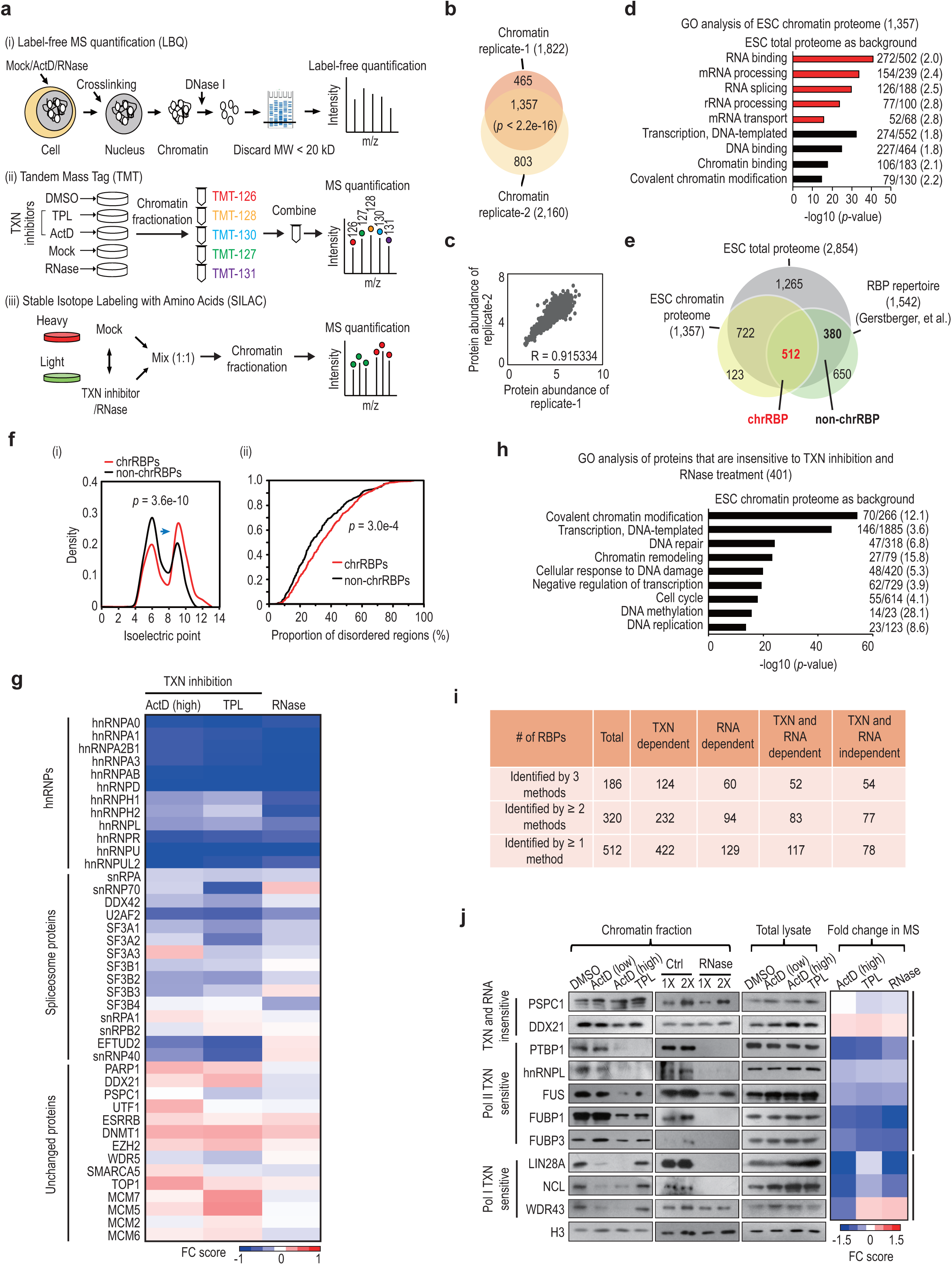
Abundant and dynamic associations of RBPs with chromatin in ESCs. **a,** Schemes showing quantitative analysis of chromatin proteomes under various treatments by three mass spec (MS) methods. (i) Label-free MS quantification (LBQ); (ii) Tandem mass tag (TMT); (iii) Stable isotope labeling with amino acids (SILAC). Transcription (TXN) inhibition: ActD (actinomycin D, 1 μg/ml) or TPL (triptolide, 1 μM). RNase: RNase A (1 mg/ml). DMSO/Mock: mock treatment for transcription inhibition or RNase treatment. **b,** Overlap between two biological replicates of the chromatin proteome. *P*-values, Fisher’s exact test. **c,** Correlation analysis of two biological replicates of the chromatin proteome. The x-axis and y-axis represent the abundance of each protein identified in the two replicates, indicated by −log10 (iBaq ratio) (see materials and methods). See also Supplementary Table 1. **d,** Gene ontology (GO) analysis of chromatin proteins (n = 1,357). The total proteome from ESCs (n = 2,854) was used as background. Selected GO terms (*p* < 1.0e-10) are shown on the y-axis. The x-axis shows enrichment significance by −log10 (*p*-value). Red bars represent terms related to RNA processes; black bars indicate terms associated with transcription and chromatin functions. For each GO term, the number of functionally associated genes identified from analysis of the chromatin proteome and the total number of functionally associated genes expressed in ESCs are indicated sequentially. The numbers in the brackets indicate the fold enrichment. **e,** Comparison of the chromatin proteome (n = 1,357) with the RBP repertoire (n = 1,542) and the ESC total proteome (n = 2,854). The numbers of chrRBPs (red, 512) and non-chrRBPs (black, 380) are indicated in bold. See also Supplementary Table 2. **f,** Biochemical characterization of chrRBPs and non-chrRBPs. (i) Density distribution curve of the isoelectric points of chrRBPs and non-chrRBPs. The blue arrow indicates a shift in the distribution of isoelectric point. (ii) Cumulative distribution curve showing the content of intrinsically disordered regions (IDR) in chrRBPs or non-chrRBPs. *P*-values, Kolmogorov-Smirnov test. See also Supplementary Table 2. **g,** Heatmap showing the average fold change (FC) score of chromatin abundance for representative proteins including hnRNPs, spliceosome proteins and unchanged proteins. Numerous hnRNPs and splicing factors are dependent on both RNA and transcription for their chromatin binding. In comparison, the chromatin-binding activities of transcription factors (such as UTF1 and ESRRB) and epigenetic enzymes (such as DNMT1, EZH2, WDR5, topoisomerases, and DNA helicases) were less likely to be affected. The ratio calculation is described in Materials and Methods. Data are shown as the mean of 4 biological replicates for ActD and RNase, and 3 replicates for TPL. See also Supplementary Table 3. **h,** GO analysis of chromatin proteins that are insensitive to transcription inhibition and RNase treatment (n = 401). The ESC chromatin proteome was used as the background. The x-axis shows enrichment significance by −log10 (*p*-value). The top enriched terms are shown on the y-axis. For each GO term, the number of functionally associated genes identified from analysis of the chromatin proteome and the total number of functionally associated genes expressed in ESCs are indicated sequentially. The numbers in the brackets indicate the fold enrichment. **i,** Summary of the effects of transcription (TXN) and RNA on chromatin-RBP associations. See also Supplementary Table 3. **j,** Chromatin fraction and western-blot analysis of selected chrRBPs upon treatments with transcription inhibitors or RNase A. The corresponding FC score in the mass spec data is shown in heatmap (right). ActD: low (10 ng/ml) or high (1 μg/ml). RBPs are classified into 3 groups based on their sensitivity to inhibition of Pol I or Pol II transcription (TXN) or RNA. Because nascent transcripts are loaded and protected by a battery of RBPs once they emerge from Pol II^106^, we cannot rule out incomplete degradation of RNA by treatment with RNase A. Thus, despite an overall decrease of chrRBP associations with chromatin, the role of RNA in recruiting and mediating chrRBPs to chromatin might be underestimated based on the observed effects of RNase treatment.

**Extended Data Fig. 2.**
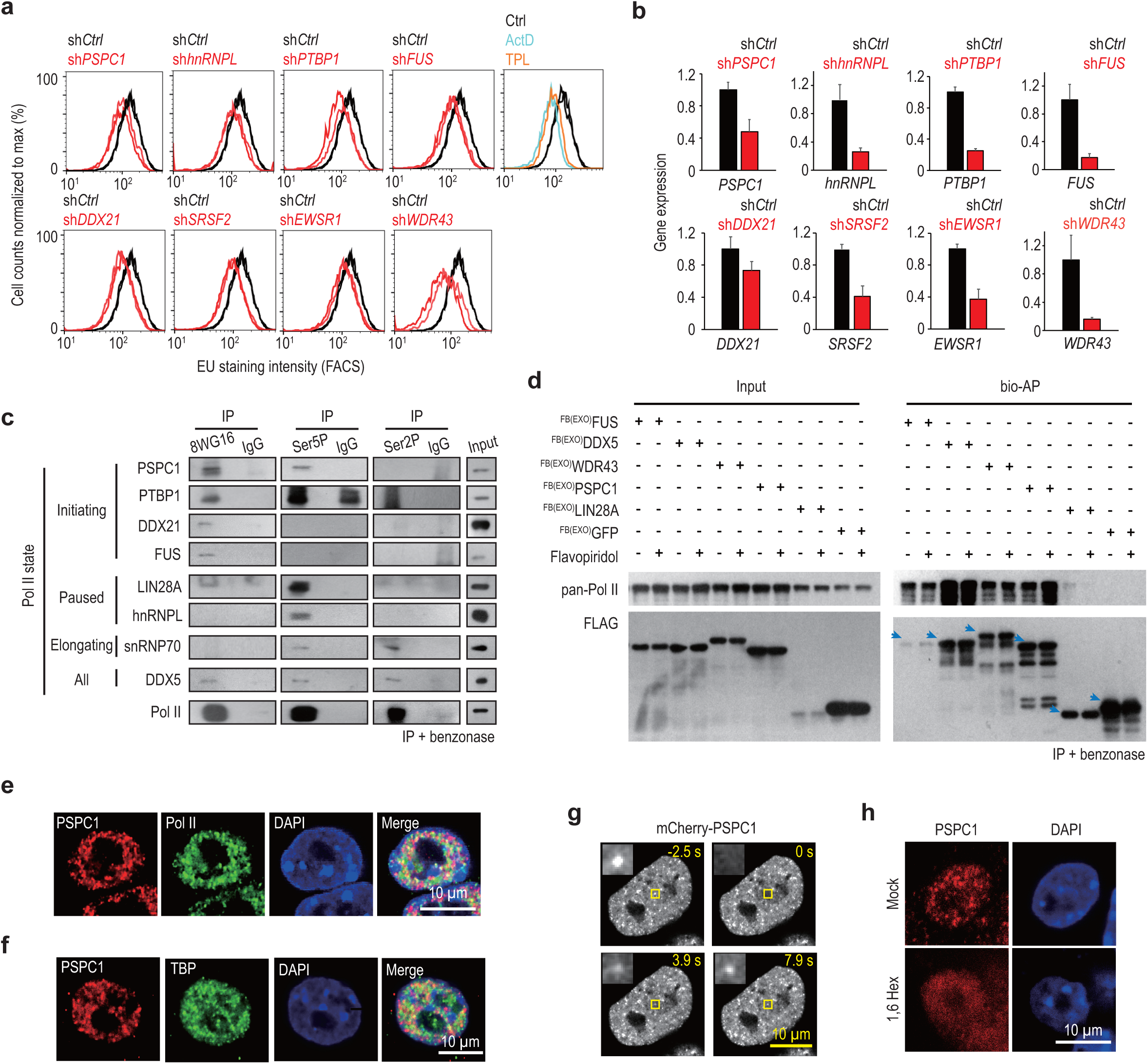
chrRBPs tend to interact with Pol II and modulate transcription. **a,** Fluorescence activated cell sorting (FACS) of 5-Ethynyl uridine (EU) incorporation. The x-axis shows EU intensity. The y-axis shows cell numbers normalized to max (%). Two biological replicates are shown for controls (black) and RBP knockdown (red). Depletion of individual chrRBPs all led to reduced EU staining of newly synthesized transcripts, to a degree slightly weaker, yet comparable to that caused by knockdown of WDR43, a critical regulator of Pol II pause release and Pol I transcription^77^, or by treatments with the transcription inhibitors ActD and TPL. **b,** RT-qPCR analysis of the relative expression of RBPs upon knockdown for 60 hours. Data are shown as mean ± s.d. of 2 biological replicates. **c,** Co-immunoprecipitation (co-IP) and western blot analysis of endogenous proteins. Pol II in different phosphorylation states captured all 8 tested chrRBPs in benzonase-treated native ESC lysates, indicating RNA/DNA-independent interactions. For example, endogenous PSPC1 was pulled down by initiating (hypoP) and paused (Ser5P) Pol II, but not by elongating Pol II (Ser2P). Capture of various phosphorylation states of Pol II was confirmed by western blots using the corresponding antibody in each IP (bottom). Benzonase was added during cell lysis and co-IP. 8WG16, hypo-phosphorylated (hypoP) Pol II that represents initiating Pol II. Ser5P, serine 5-phosphorylated Pol II that represents paused Pol II. Ser2P, serine 2-phosphorylated Pol II that represents elongating Pol II^107^. **d,** Reciprocal co-IP of various chrRBPs captured Pol II. Biotin-mediated affinity purification (bio-AP) was performed (+ benzonase) in ESCs that stably express individual FLAG-biotin tagged chrRBPs. ESCs expressing FLAG-biotin tagged GFP (^FB(EXO)^GFP) were used as the negative control. Flavopiridol was used to inhibit transcription. PSPC1, FUS, DDX5, and WDR43 captured pan-Pol II, independently of transcription and/or DNA/RNA, whereas LIN28A exhibited weak interaction with Pol II in a transcription-dependent manner. Blue arrows indicate exogenously expressed FLAG-tagged proteins. **e,** Co-immunostaining analysis of PSPC1 with pan-Pol II. The experiment was performed in ^AID-FB(KI)^*PSPC1* cells and anti-FLAG antibody was used to image endogenously tagged PSPC1 proteins. **f,** Co-immunostaining analysis of PSPC1 with TBP. This experiment was performed in wild-type ESCs and PSPC1 antibody was used to detect endogenous proteins. **g,** Fluorescence recovery after photobleaching (FRAP) analysis showing the fast recovery of mCherry-PSPC1 puncta. Related to Fig. 1i. **i,** Immunofluorescence analysis of PSPC1 upon treatment with 10% 1,6-hexanediol for 2 minutes. Mock: PBS. Anti-PSPC1 antibody was used for imaging.

**Extended Data Fig. 3.**
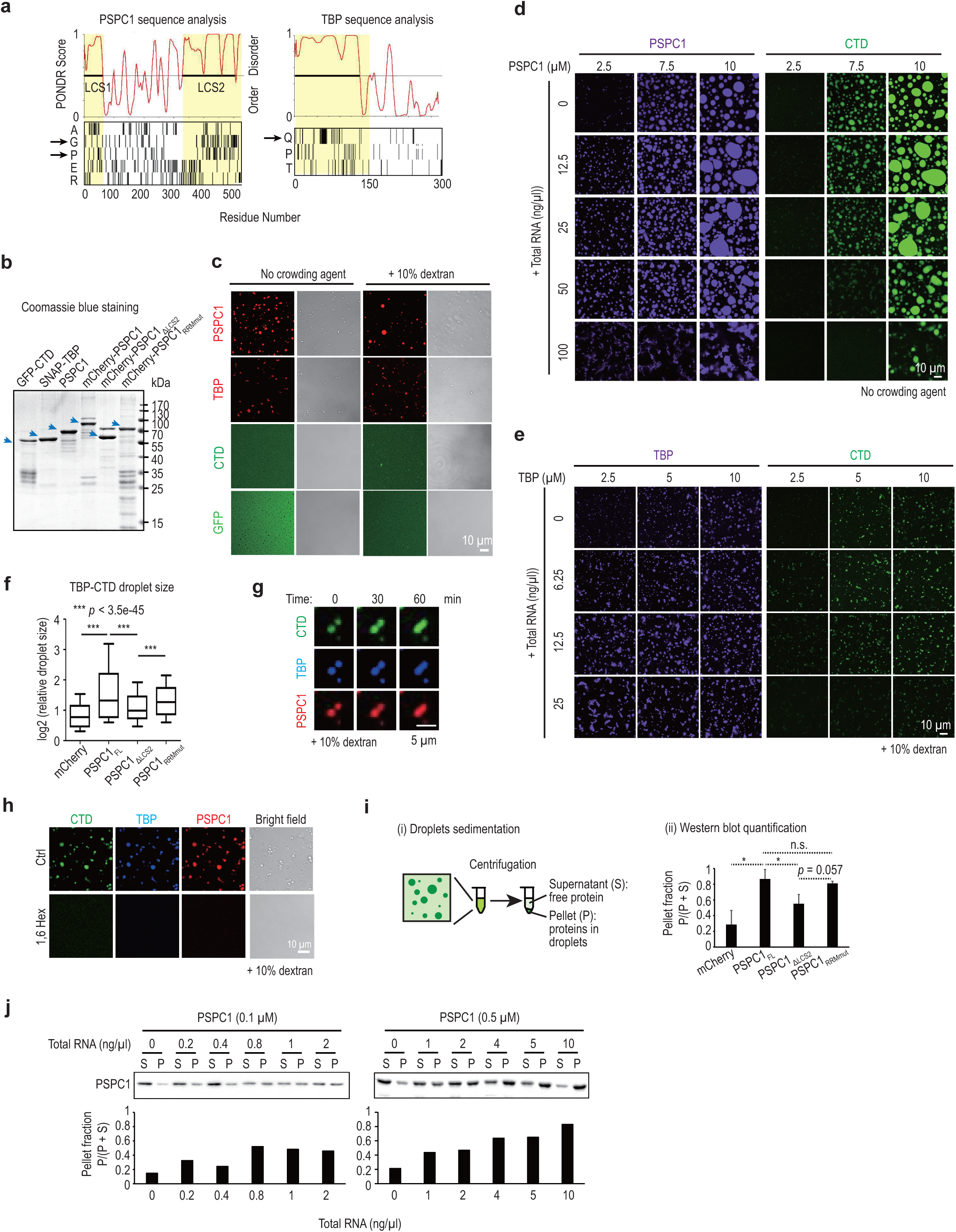
PSPC1 promotes the incorporation of unphosphorylated CTD into TBP condensates. **a,** Analysis of disordered regions and low-complexity sequences in PSPC1 and TBP. The PONDR score indicates the probability of a region being disordered. Regions with a score > 0.5 are defined as disordered. The distributions of representative amino acids (alanine, A; glycine, G; proline, P; Glutamic acid, E; arginine, R; glutamine, Q; threonine, T) are shown at the bottom. The regions in yellow indicate the disordered regions of PSPC1 and TBP which contain the hydrophobic G/P-rich or Q-rich repetitive sequences respectively (arrow). **b,** Coomassie blue staining of purified HIS-tagged GFP-CTD, SNAP tagged-TBP, PSPC1, mCherry tagged-PSPC1, mCherry tagged-PSPC1_ΔLCS2_, and mCherry tagged-PSPC1_RRMmut_. The blue arrows indicate the main band of the corresponding protein. **c,** *In vitro* droplet formation of recombinant PSPC1, TBP, CTD and GFP proteins. The assay was performed with 5 μM of proteins in a solution containing 150 mM NaCl with or without the crowding agent dextran. mCherry-PSPC1 was mixed with untagged PSPC1 (1:3). TBP was pre-stained using SNAP-647 and mixed with unlabeled protein (1:10). PSPC1 showed strong phase separation activity regardless the presence of dextran. Recombinant TBP formed fiber-like irregular aggregates in the absence of dextran, but was able to form liquid-like droplets in the presence of dextran, at a concentration of 5 μM, which is well above its estimated nuclear concentration of 0.06∼0.3 μM. Thus, we added dextran for subsequent TBP-involved droplet assays. CTD with 20 heptad repeats also failed to phase-separate at 5 μM even with 10% dextran. **d-e,** Phase diagram of PSPC1-CTD or TBP-CTD droplets in the presence of different concentrations of RNA. We used 0.6 μM of CTD in all assay conditions, and tested increasing concentrations of PSPC1 or TBP from 2.5 μM to 10 μM as indicated. PSPC1 and TBP were pre-stained using Cy5.5 or SNAP-647 and were mixed with unlabeled protein (1:10). RNA was total RNA isolated from ESCs. No crowding agent was added for PSPC1 in panel **d**, while 10% dextran was added for TBP in panel **e**. **f,** Quantification for TBP-CTD droplet size in droplet formation assays of TBP (5 μM) and CTD (0.6 μM) with full-length (FL) or mutant PSPC1 (5 μM) or mCherry (5 μM). All pictures were acquired at the same time. Representative pictures are shown in Fig. 2a. The y-axis is log2 (relative droplet size). The median droplet size is 0.71 μm^2^ for mCherry (n = 2,641), 1.50 μm^2^ for PSPC1_FL_ (n = 2,932), 0.99 μm^2^ for PSPC1_ΔLCS_ (n = 2,097), and 1.42 μm^2^ for PSPC1_RRMmut_ (n = 4,758). *P*-values, two-sided Student’s t-test. **g,** Fusion of TBP-CTD-PSPC1 droplets. Representative images are shown. TBP (5 μM), CTD (0.6 μM) and PSPC1 (5 μM) were incubated in a solution containing 150 mM NaCl and 10% dextran. **h,** Phase-separated droplets composed of TBP, CTD and PSPC1 with or without treatment by 10% of 1,6-hexanediol. TBP (5 μM), CTD (0.6 μM) and PSPC1 (5 μM) were incubated in a solution containing 150 mM NaCl and 10% dextran. Ctrl: mock treatment. **i-j,** Droplets sedimentation and western-blot assays. In panel **i**, the schematic diagram and quantification are shown in panel (i) and (ii), respectively. Representative western-blot result is shown in Fig. 2c. The pellet fraction ratio P/(S + P) was shown as mean ± s.d. of ⩾2 independent biological replicates calculated based on the quantified western-blot results. Panel **j** shows RNA’s effects on PSPC1 phase separation. The bottom panel indicates the quantification of western-blot results. When no RNA was added, only small fraction of PSPC1 (∼10% at 0.1 μM and ∼20% at 0.5 μM) was present in the pellet. With addition of total RNAs, increasing proportion of PSPC1 (up to ∼80%) appears in the pellet, indicating that RNA is a multivalent ligand to promote PSPC1 phase behaviors. No crowding agent was added.

**Extended Data Fig. 4.**
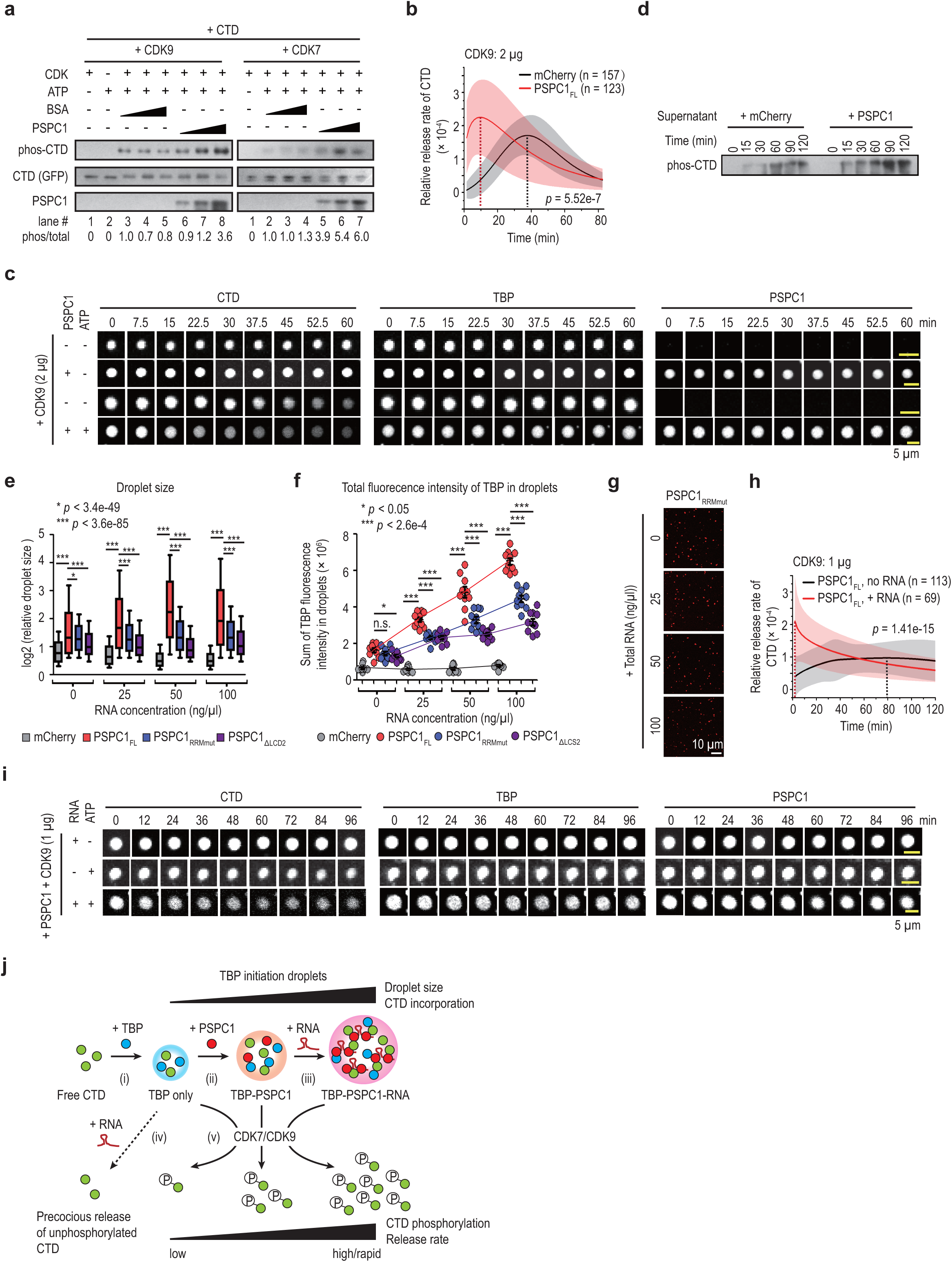
RNA synergizes with PSPC1 in promoting Pol II incorporation, phosphorylation and release. **a,** Western-blot analysis of kinase assays with CDK9 (left) and CDK7 (right). Increasing amounts (1-5 μg) of PSPC1 or BSA were incubated with CTD (0.2 μg). CDK9/CDK7 (0.2 μg) and ATP (final 0.1 mM) were added into the reaction. The antibody used for the detection of phos-CTD is anti-Pol II Ser5P. Quantification of the ratio of phosphorylated CTD versus total CTD is shown at the bottom. **b-c,** Time-lapse imaging analysis of CTD release. CDK9 (2 μg) and ATP (final 0.1 mM) were added to phase-separated droplets composed of TBP (5 μM), CTD (0.6 μM) and PSPC1 or mCherry (5 μM) to initiate the reaction. Panel **b** shows the relative release rate of CTD (y-axis). The rate calculation was described in Materials and Methods. Data are shown as mean ± s.d. of 157 droplets for the mCherry group and 123 droplets for the PSPC1_FL_ group. *P*-value, two-tailed Student’s t-test for the comparison of max rate between the two groups. Panel **c** shows images taken of representative droplets under each condition. The CTD channel (left), TBP channel (middle) and PSPC1 channel (right) of individual droplets were recorded simultaneously. Related to Fig. 2e-f. **d,** Time-lapse sedimentation and western-blot analysis of released CTD. CDK9 (2 μg) and ATP (final 0.1 mM) were added to phase-separated droplets composed of TBP (5 μM), CTD (0.6 μM) and PSPC1 or mCherry (5 μM) to initiate the reaction. At each indicated time point, droplets and free protein were collected by sedimentation. The same fraction of supernatant from each reaction (PSPC1 or mCherry) was loaded for western-blot analysis. The samples are from the same experiments in Fig. 2g. The antibody used for the detection of phos-CTD is anti-Pol II Ser5P. **e,** Effects of RNA on CTD incorporation into TBP droplets in the presence or absence of PSPC1 proteins. TBP (5 μM), CTD (0.6 μM), various PSPC1 proteins (5 μM) and mCherry (5 μM) were used in the assay. Quantification of droplet size was based on images obtained in Fig. 2h. The droplet sizes are presented relatively as log2 (relative droplet size). The median droplet sizes measured as surface area are shown in the sequence of ‘no RNA’, ‘25 ng/μl RNA’, ‘50 ng/μl RNA’, and ‘100 ng/μl RNA’. mCherry group: 0.71 μm^2^ (n = 2,641), 0.56 μm^2^ (n = 2,555), 0.40 μm^2^ (n = 1,698), and 0.40 μm^2^ (n = 1,961). PSPC1_FL_ group: 1.50 μm^2^ (n = 2,932), 2.19 μm^2^ (n = 2,732), 3.69 μm^2^ (n = 1,838), and 2.79 μm^2^ (n = 3,663). PSPC1_RRMmut_ group: 1.42 μm^2^ (n = 4,758), 1.37 μm^2^ (n = 5,029), 1.50 μm^2^ (n = 7,073), and 1.50 μm^2^ (n = 8,567). PSPC1_ΔLCD2_ group: 0.99 μm^2^ (n = 2,097), 0.94 μm^2^ (n = 1,313), 0.82 μm^2^ (n = 1,436), and 1.03 μm^2^ (n = 1,429). *P*-values, two-sided Student’s t-test. n.s., not significant. **f,** Quantifications of total fluorescence intensity of TBP in droplets shown in Fig. 2h. N = 10 fields for each condition. *P*-values, two-sided Student’s t-test. n.s., not significant. **g,** Effects of RNA on PSPC1_RRMmut_ droplets. PSPC1_RRMmut_ protein (5 μM) was incubated in a solution containing 150 mM NaCl and increasing concentration of RNA (0-100 ng/μl). No significant changes were observed, indicating the significant roles of RNA-binding abilities in promoting PSPC1 phase separation. **h-i,** Time-lapse imaging analysis of CTD release with or without RNA. CDK9 (1 μg) and ATP (final 0.1 mM) were added to phase-separated droplets composed of TBP (5 μM), CTD (0.6 μM) and PSPC1 (5 μM) in the presence or absence of RNA (50 ng/μl). Because CTD signals declined sharply from TBP condensates in the presence of RNA and PSPC1 immediately after addition of CDK9 and ATP, we slowed down the kinase reaction by adding less CDK9 enzyme (1 μg) in order to monitor the time course of CTD release, as compared to Extended Data Fig. 4b-c. Panel **g** shows the quantification of CTD release rate. Data are shown as mean ± s.d. of 113 droplets for the ‘PSPC1_FL_, no RNA group’ and 69 droplets for the ‘PSPC1_FL_, + RNA’ group. *P*-value, two-tailed Student’s t-test for the comparison of max rate between the two groups. Panel **i** shows images taken of representative droplets under each condition. The CTD channel (left), TBP channel (middle) and PSPC1 channel (right) were recorded simultaneously for individual droplets. Related to Fig. 2k-l. **j,** Schematic diagram showing the interplay of PSPC1 and RNA in promoting CTD incorporation and subsequent phosphorylation and release. TBP alone has weak ability to phase separate and trap CTD within its droplets (i). Addition of PSPC1 enhances this phase separation and produces larger droplets that concentrate more CTD inside (ii). RNA further synergizes with PSPC1 to drastically promote phase separation and CTD incorporation (iii). By contrast, in the absence of PSPC1, RNA evicts CTD from TBP droplets (iv). Upon activation by CTD kinases (v), efficient compartmentalization and concentration of CTD inside TBP-PSPC1-RNA droplets lead to stronger phosphorylation and faster release of CTD compared to TBP and TBP-PSPC1 droplets. Note that RNA synergizes with PSPC1 in a manner that critically depends on the phase-separation and RNA-binding activities of PSPC1. Thus, *in vitro* assays with defined components allow us to biochemically dissect the more complex processes of Pol II engagement and release in cells.

**Extended Data Fig. 5.**
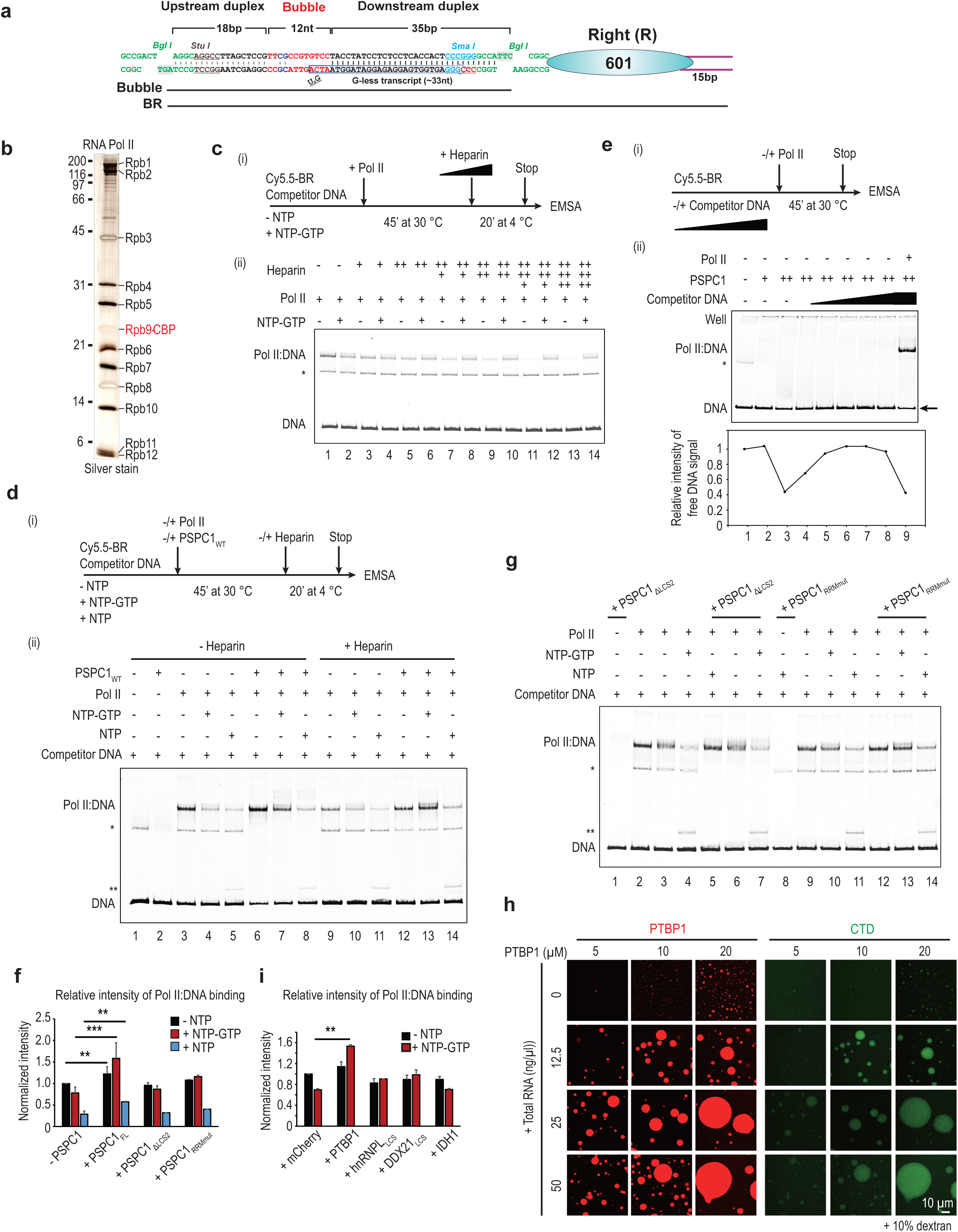
PSPC1 promotes and stabilizes Pol II binding during transcription *in vitro*. **a,** Schematic diagram of the template DNA. ‘BR’ stands for bubble right (the 601 sequence). The BR template was labeled with either biotin or Cy5.5. **b,** Silver staining of purified Pol II from yeast. **c,** Titration of heparin in the EMSA assay to reduce unspecific docking of Pol II on the BR template. The concentration of heparin used in lanes 3-4 was chosen for *in vitro* transcription and EMSA assays. **d,** EMSA of Pol II and BR template during *in vitro* transcription. The BR template (Cy5.5-labeled), Pol II, and PSPC1 in the absence or presence of NTPs as indicated were incubated at 30 °C for 45 minutes. Heparin was then added and incubated at 4 °C for 20 minutes to reduce the non-specific binding of Pol II. The free template (‘DNA’) and the supershifted ‘Pol II:DNA’ bands are indicated on the left. The bands marked by single asterisk is likely to be a non-specified byproduct during BR template assembly and gel purification. The bands marked by double asterisks is likely to be a R-loop, given its sensitivity to RNase H (data not shown). Heparin effectively removed the docking Pol II from the template in the absence of NTPs (comparing lane 9 to lane 3), but had negligible effects on the stalled or elongating Pol II (comparing lane 10 to 4 and lane 11 to 5). Importantly, addition of recombinant PSPC1_FL_ consistently enhanced the Pol II:DNA signals in both the absence (lane 3-5 vs lane 6-8) and the presence of heparin (lane 9-11 vs 12-14). **e,** Titration of a competitor DNA to reduce nonspecific binding of PSPC1 to the BR template. Quantification of the free template signal (indicated as ‘DNA’ by an arrowhead) was shown at the bottom. In the absence of a competitor DNA, PSPC1 exhibited a weak binding affinity to the template, as addition of PSPC1 decreased the amount of free template DNA signals and increased DNA signals stuck in the well (comparing lanes 2-3 to lane 1). Upon addition of the competitor DNA, more free DNA signals were detected (lanes 4-8), which suggests that less DNA template was bound by PSPC1. The amount of competitor DNA used in lanes 8-9 was chosen for *in vitro* transcription and EMSA assays. In this condition, the binding of PSPC1 to template DNA is minimized, while Pol II’s binding was not affected (lane 9). **f,** Summary of Pol II:DNA signals in several independent experiments shown in Fig. 3c-d, Extended Data Fig. 5e and 5g, and in biological replicates not shown here. Only reactions with the addition of heparin were quantified. The y-axis shows the band intensity normalized to the reaction without addition of PSPC1. Data are shown as mean ± s.d. of ≥ 2 biological replicates for each condition. *P*-values, two-sided Student’s t-test. **g,** Effects of PSPC1 mutants on the binding of Pol II to BR template during *in vitro* transcription (heparin included). PSPC1_RRMmut_ and PSPC1_ΔLCS2_ had negligible effects on Pol II binding (lane 5-7 and 12-14; Extended Data Fig. 5f). **h,** Phase diagram of PTBP1-CTD droplets in the presence of different concentrations of RNA. We used 0.6 μM of CTD in all assay conditions, and tested increasing concentrations of PTBP1 from 5 μM to 20 μM as indicated. RNA was total RNA isolated from ESCs. 10% dextran was added. **i,** Summary of Pol II:DNA signals in several independent experiments shown in Fig. 3e and in biological replicates not shown here. Data are shown as mean ± s.d. of ≥ 2 biological replicates for each condition. *P*-values, two-sided Student’s t-test.

**Extended Data Fig. 6.**
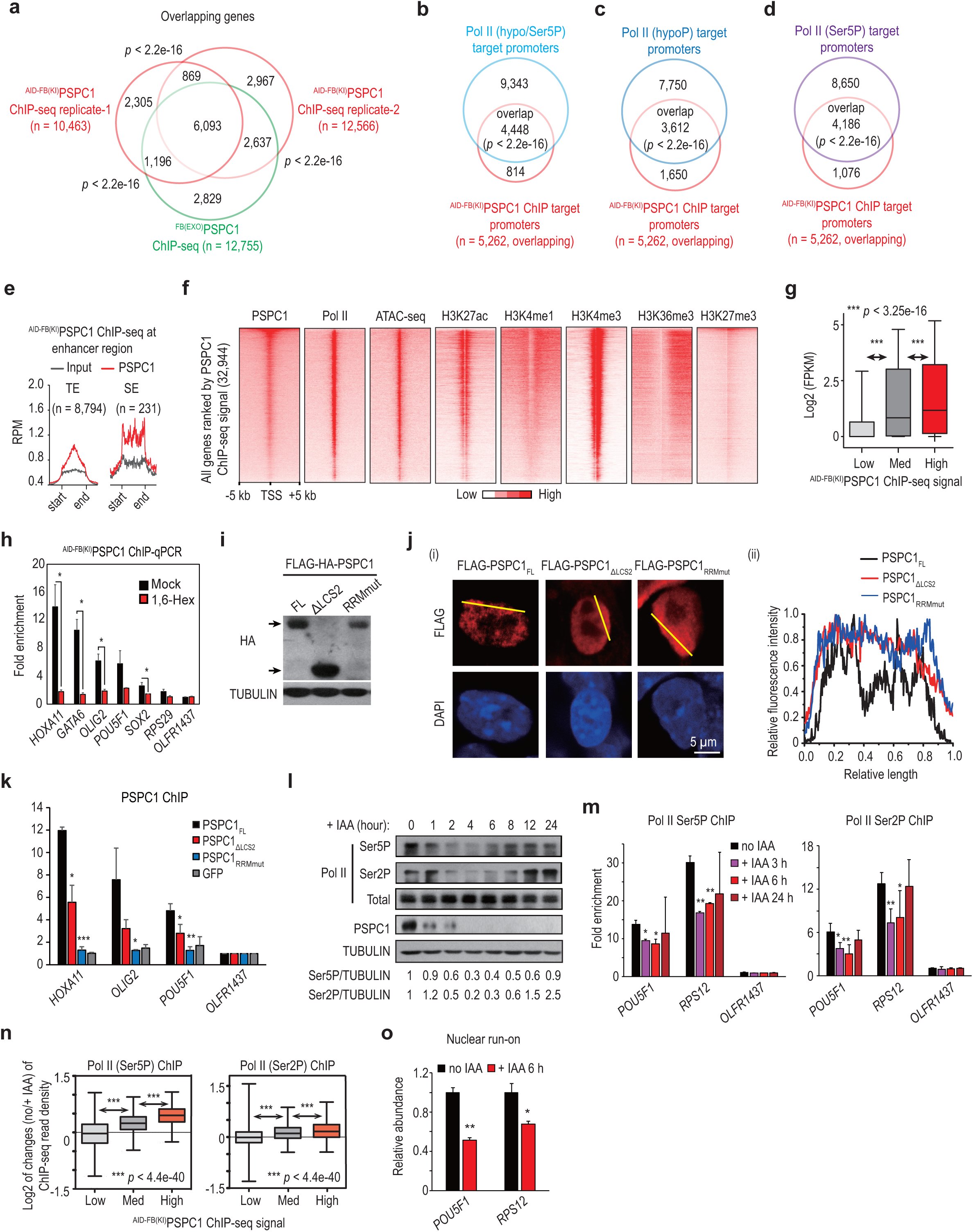
Time-course analysis of PSPC1 degradation by AID. **a-g,** ChIP-seq analysis of PSPC1. Panel **a** shows overlap between targets identified by two biological replicates of ^AID-FB(KI)^PSPC1 ChIP-seq and one ^FB(EXO)^PSPC1 ChIP-seq experiment. See also Supplementary Table 4. Panel **b** shows the overlap of ^AID-FB(KI)^PSPC1-targeted promoters (n = 5,262, overlapping promoters of two-biological replicates) and Pol II (hypo/Ser5P)-targeted promoters (13,791). Panel **c-d** shows the overlap of target promoters between ^AID-FB(KI)^PSPC1 and Pol II (hypoP) **(c)** or Pol II (Ser5P) ChIP-seq **(d)** respectively. *P*-values for panels (**a-d**) were all determined by Fisher’s exact test. Panel **e** shows metagene analysis of ChIP-seq signals of ^AID-FB(KI)^PSPC1 across enhancers. The y-axis is reads per million reads (RPM). TE, typical enhancers (n = 8,704). SE, super enhancers (n = 231). Panel **f** shows heatmaps of ChIP-seq signals of PSPC1, hypoP Pol II, histone marks, and ATAC-seq signals around TSS (± 5 kb) across all mouse genes (n = 32,944). The heatmap is sorted by PSPC1 ChIP-seq signal. Panel **g** shows the correlation between gene expression and PSPC1 ChIP-seq signals. All genes (n = 32,944) are classified equally into three groups according to PSPC1 ChIP-seq signal. The y-axis is log2 (FPKM). *P*-values, two-sided Student’s t-test. **h,** ChIP-qPCR analysis of ^AID-FB(KI)^PSPC1 with or without treatment with 1.5% of 1,6-hexanediol for 30 minutes. The relative fold enrichment at each target was normalized to an untargeted gene *OLFR1437*. Data are shown as mean ± s.d. of 2 biological replicates. *, *p* < 0.05 by two-sided Student’s t-test. **i,** Western-blot analysis of ectopically expressed FLAG-HA-tagged PSPC1 proteins in ^AID-FB(KI)^PSPC1 cells. **j,** Immunofluorescence analysis of wild-type and mutant PSPC1 proteins. Various FLAG-tagged PSPC1 proteins were transiently expressed in ESCs and were imaged by the anti-FLAG antibody at 48 hours post-transfection. Representative images are shown in (i). Relative fluorescence intensities along the yellow lines are shown in (ii). **k,** Anti-FLAG ChIP-qPCR analysis of the full-length (FL) and mutant proteins of PSPC1 that are transiently expressed in wild-type ESCs. The relative fold enrichment at each target was normalized to an untargeted gene *OLFR1437*. Data are shown as mean ± s.d. of ≥2 biological replicates. *, *p* < 0.05, **, *p* < 0.01, ***, *p* < 0.001 by two-sided Student’s t-test. **l,** Time-course western-blot analysis of Pol II and PSPC1 levels in ^AID-FB(KI)^PSPC1 cells treated with IAA. The relative levels of Ser5P and Ser2P normalized to TUBULIN are shown at the bottom. Degradation of PSPC1 dramatically altered the levels of phosphorylated Pol II, but not total Pol II. Through a 24-hour time course, we observed an initial downregulation of both Ser5P and Ser2P Pol II, with 20-30% remaining at 4 hours. Afterwards, their levels gradually increased and returned close to the original level at 24 hours, implying the existence of compensatory mechanisms that safeguard steady-state activities of Pol II. **m,** ChIP-qPCR analysis of Pol II Ser5P (left) and Ser2P (right) upon PSPC1 degradation for 3 h to 24 h. The y-axis indicates the relative fold enrichment normalized to the non-targeted gene *OLFR1437*. Data are shown as mean ± s.d. of ≥ 3 biological replicates. *, *p* < 0.05; **, *p* < 0.01 by two-sided Student’s t-test. **n,** Correlation analysis between changes in Pol II ChIP-seq and PSPC1 ChIP-seq signals. All genes (n = 32,944) are classified equally into three groups according to PSPC1 ChIP-seq signals. The y-axis shows log2 of the average change of Pol II ChIP-seq signals. *P*-values, two-sided Student’s t-test. **o,** Nuclear run-on analysis upon degradation of PSPC1 for 6 h. The y-axis indicates the relative abundance of nascent transcripts calculated by normalizing to the expression of mature *ACTB* transcript. Data are shown as mean ± s.d. of 2 biological replicates. *, *p* < 0.05; **, *p* < 0.01 by two-sided Student’s t-test.

**Extended Data Fig. 7.**
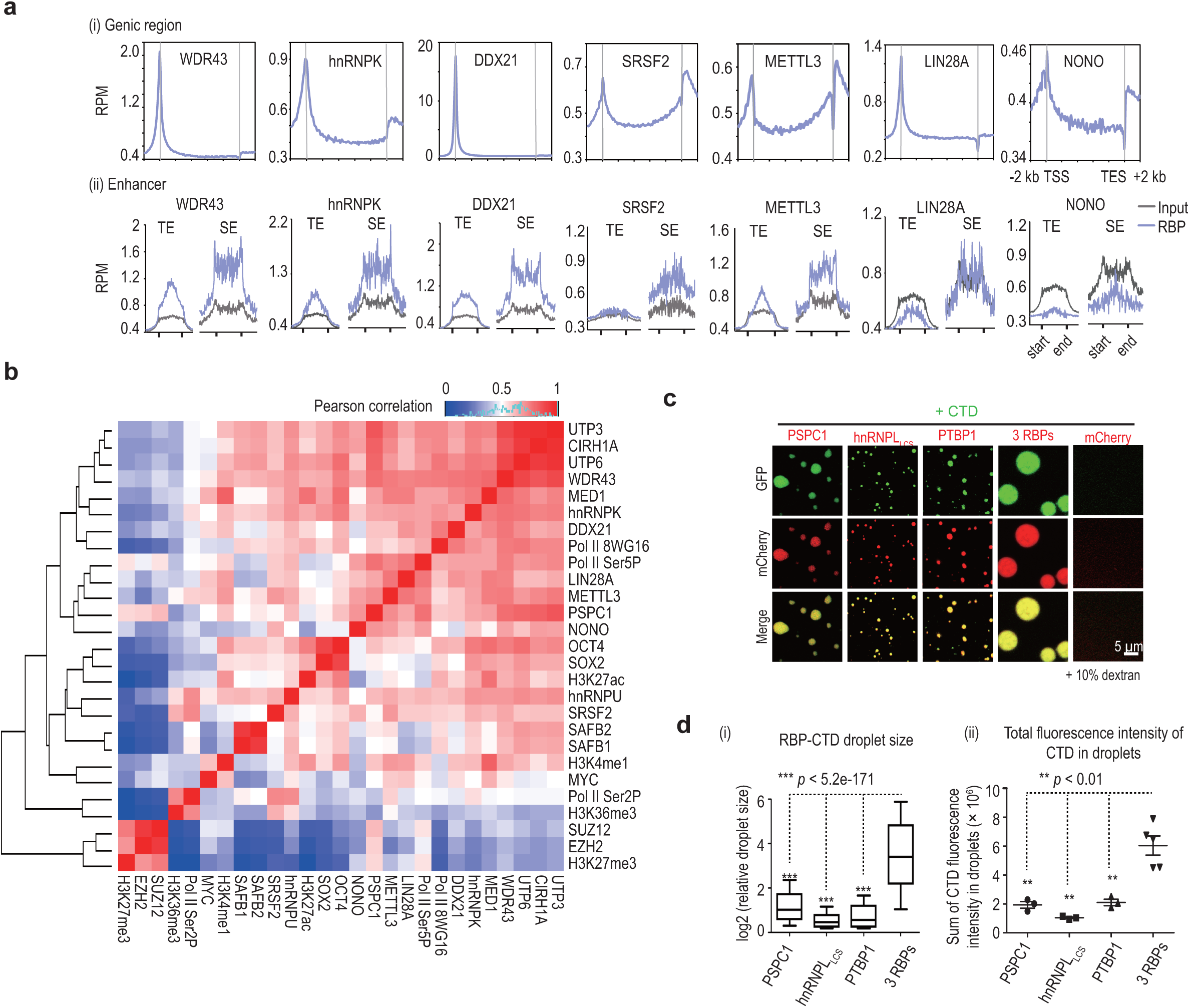
Genome-wide co-occupancy of chrRBPs with Pol II at promoters and enhancers. **a,** Metagene analysis of ChIP-seq signals of various RBPs across all mouse genes (n = 32,944) (i) and enhancers (ii). The y-axis is reads per million reads (RPM). TE, typical enhancers (n = 8,704). SE, super enhancers (n = 231). Related to Fig. 5a. **b,** Heatmap showing hierarchical clustering of chromatin binding by RBPs, by transcription regulators, and histone modifications in ESCs. The color indicates the Pearson correlation value. **c-d,** Phase-separation assay of various RBPs with the CTD. Compared to mCherry-tagged PSPC1 (5 μM), mCherry-tagged PTBP1 (10 μM) and mCherry-tagged LCS domain of hnRNPL (hnRNPL_LCS_, 30 μM) exhibited weak phase-separation activity and incorporated GFP-tagged CTD (0.6 μM) inside their droplets. mCherry with the highest concentration that equals the sum of all proteins (45 μM) was used as a control, which remains clear. The assay was performed in 150 mM NaCl and 10% dextran. Representative pictures are shown in panel **c**. Quantification for RBP-CTD droplet size (i) and the total CTD fluorescence intensity (ii) in the droplet are summarized in panel **d**. The y-axis is log2 (relative droplet size) in the upper panel. The median droplet size is 1.04 μm^2^ for PSPC1 (n = 782), 0.37 μm^2^ for hnRNPL_LCS_ (n = 6,158), 0.48 μm^2^ for PTBP1 (n = 2,374), and 9.60 μm^2^ for all 3 RBPs together (n = 554). In the bottom panel, the y-axis shows the sum of fluorescence intensity of CTD in droplets in each field of view. N ⩾ 3 fields for each condition. *P*-values, two-sided Student’s t-test.

